# Choice of host cell line is essential for the functional glycosylation of the fragment crystallizable (Fc) region of human IgG1 inhibitors of influenza B viruses

**DOI:** 10.1101/719849

**Authors:** Patricia A. Blundell, Dongli Lu, Anne Dell, Stuart M. Haslam, Richard J. Pleass

**Author notes:** To whom correspondence should be addressed: Dept. of Parasitology, Liverpool School of Tropical Medicine, Pembroke Place, Liverpool, L3 5QA, United Kingdom.

## Abstract

Antibodies are glycoproteins that carry a conserved N-linked carbohydrate attached to the Fc, whose presence and fine structure profoundly impacts on their *in vivo* immunogenicity, pharmacokinetics and functional attributes. The host cell line used to produce IgG has a major impact on this glycosylation, as different systems express different glycosylation enzymes and transporters that contribute to the specificity and heterogeneity of the final IgG-Fc glycosylation profile. Here we compare two panels of glycan-adapted IgG1-Fc mutants expressed in either the HEK 293-F or CHO-K1 systems. We show that the types of N-linked glycans between matched pairs of Fc mutants vary significantly, and in particular with respect to sialylation. These cell line effects on glycosylation profoundly influence the ability of the engineered Fcs to interact with either human or pathogen receptors. For example, we describe Fc mutants that potently disrupted influenza B-mediated agglutination of human erythrocytes when expressed in CHO-K1 but not in HEK 293-F cells.

## Introduction

In therapeutic approaches where the Fc of human IgG1 is critically important, receptor binding and functional properties of the Fc are lost after de-glycosylation or removal of the Asn-297 N-linked glycosylation attachment site located in the body of the Fc (1–3). More detailed studies into the types of sugars involved in this functionality have shown enhanced FcγRIIIA binding and ADCC of IgG1 in the absence of fucose (4, 5); enhanced FcγRIIIA binding but rapid clearance from the circulation of IgG1 enriched for oligomannose structures (6–8); improved solubility, anti-inflammatory activity, thermal stability and circulatory half-life of terminally sialylated glycans from IgG1 (9–13).

These findings have generated an incentive to modify the existing IgG1 glycans attached to Asn-297, either by chemical means (12, 14), by mutagenesis programs on the Fc protein backbone that disrupt the protein-Asn-297-carbohydrate interface (15), or by expression in glycosidase-adapted transgenic cell lines (reviewed in 16). For example, the FDA approved humanized antibody Mogamulizumab, which is used to treat lymphoma and is manufactured in CHO cell lines in which the α(1-6)-fucosyltransferase (*FUT8)* gene is removed, results in an afucosylated IgG1 with enhanced FcγRIIIA-dependent tumour cell killing by ADCC (17). Although similar approaches have yielded enhanced sialylation of IgG, with zero to moderate improvements in binding to FcγRs (12, 15, 18, 19), these have not led to significant enhancements in binding to inhibitory glycan receptors that are important in controlling unwanted inflammation (19, 20), a finding we and others have attributed to the buried location of the Asn-297 attached glycan within the Fc (21, 22).

We took an alternative approach to enhancing the sialic acid content of the Fc of IgG1 (23, 24), by adding the 18 amino-acid tailpiece (tp) from IgM to the C-terminus of the IgG1 Fc, into which a cysteine-to-alanine substitution is made at Cys-575, and including an extra N-glycosylation site to the N-terminus at position Asn-221. The tp also contains a N-glycosylation site at Asn-563. When expressed in CHO-K1 cells, these molecules displayed enhanced binding to the low-affinity Fcγ-receptors (FcγRIIIA and FcγRIIB), and to multiple glycan receptors that control excessive inflammation by IVIG (23–25). Two such hyper-sialylated molecules (D221N/C575A and D221N/C309L/N297A/C575A) also bound recombinant hemagglutinin from influenza A and B viruses, and disrupted influenza A-mediated agglutination of human erythrocytes (24).

Chinese hamster ovary (CHO) cell-based systems remain by far the most common mammalian cell line used by the pharmaceutical industry; 84% of products are produced in this cell system, and the remaining approved antibodies are produced in either NS0 or Sp2/0 cells (26). Although CHO cells account for the largest number of FDA approved bio-therapeutics (26), they do not express α1,2/3/4 fucosyltransferase and β-1-4-N-acetylglucosaminyl-transferase III, which are enzymes expressed in human cells (27). Furthermore, humans have active α2,6-sialyltransferase. As such, CHO derived IgG1 Fcs are only sialylated through α2,3 linkages whereas both α2,3 and α2,6 linkages can be found on human IgG1 Fc (23, 27). Most non-human mammalian cell lines can also attach Neu5Gc. Humans do not have an active CMP-Neu5Ac hydroxylase so do not attach Neu5Gc, which can elicit immunogenic responses (27) and consequently non-human cell lines are stringently screened to identify clones that produce proteins with desirable glycan profiles (28).

Human cell lines are a promising alternative to non-human cell lines as they possess fully human post-translational modifications that reduce downstream processing costs and, more importantly, circumvent any risks associated with immunogenicity from non-human glycans. However, human cell lines also have significant limitations, including the capacity to produce sialyl-Lewis^x^ which binds to endothelial selectins in areas of inflammation (29). Although this may potentially be favourable for anti-inflammatory therapies (29), the attached sialyl-Lewis^x^ may also adversely affect the biodistribution and pharmacokinetics of a Fc when used in other clinical contexts, for example anti-tumor mAbs. Human cell lines also carry the risk of contamination and forward transmission of human pathogens, in particular viruses, that may explain why CHO-K1 cells are still the preferred cell line used by the pharmaceutical industry. These issues led us to compare the functional properties of a panel of Fc mutants generated in CHO-K1 cells with the same set of proteins manufactured by HEK 293-F cells (24).

## Materials and Methods

### Production of mutants

The generation of glycan mutants in all combinations has been described previously for the hexa-Fc that contains cysteines at both positions 309 and 575 (23). To make the new mutants described in Fig. 1 in which Cys-575 was mutated to alanine, PCR overlap extension mutagenesis was used with a pair of internal mismatched primers 5’-ACCCTGCTTGCTCAACTCT-3’ / 3’-GGCCAGCTAGCTCAGTAGGCGGTGCCAGC-5’ for each plasmid vector coding for a designated glycan modification. The parental plasmids used for these new PCR reactions have been described previously (23). The resulting C575A mutants were then further modified to remove Cys-309 using primer pair 5’-TCACCGTCTTGCACCAGGACT-3’ / 3’-AGTCCTGGTGCAAGACGGTGA-5’ to create the panel of double cysteine knockouts described in Fig. 2. To verify incorporation of the desired mutation and to check for PCR-induced errors, the open reading frames of the new mutants were sequenced on both strands using previously described flanking primers (23). CHO-K1 cells (European Collection of Cell Cultures) were stably transfected with plasmids using FuGene (Promega), stable cell lines were created, and Fc-secreting clones were expanded and proteins purified as previously described (30, 31). HEK 293-F cells were transiently transfected using the FreeStyle MAX293 expression system (Life Technologies) and proteins purified as for CHO-K1 cells.

**FIGURE 1.**
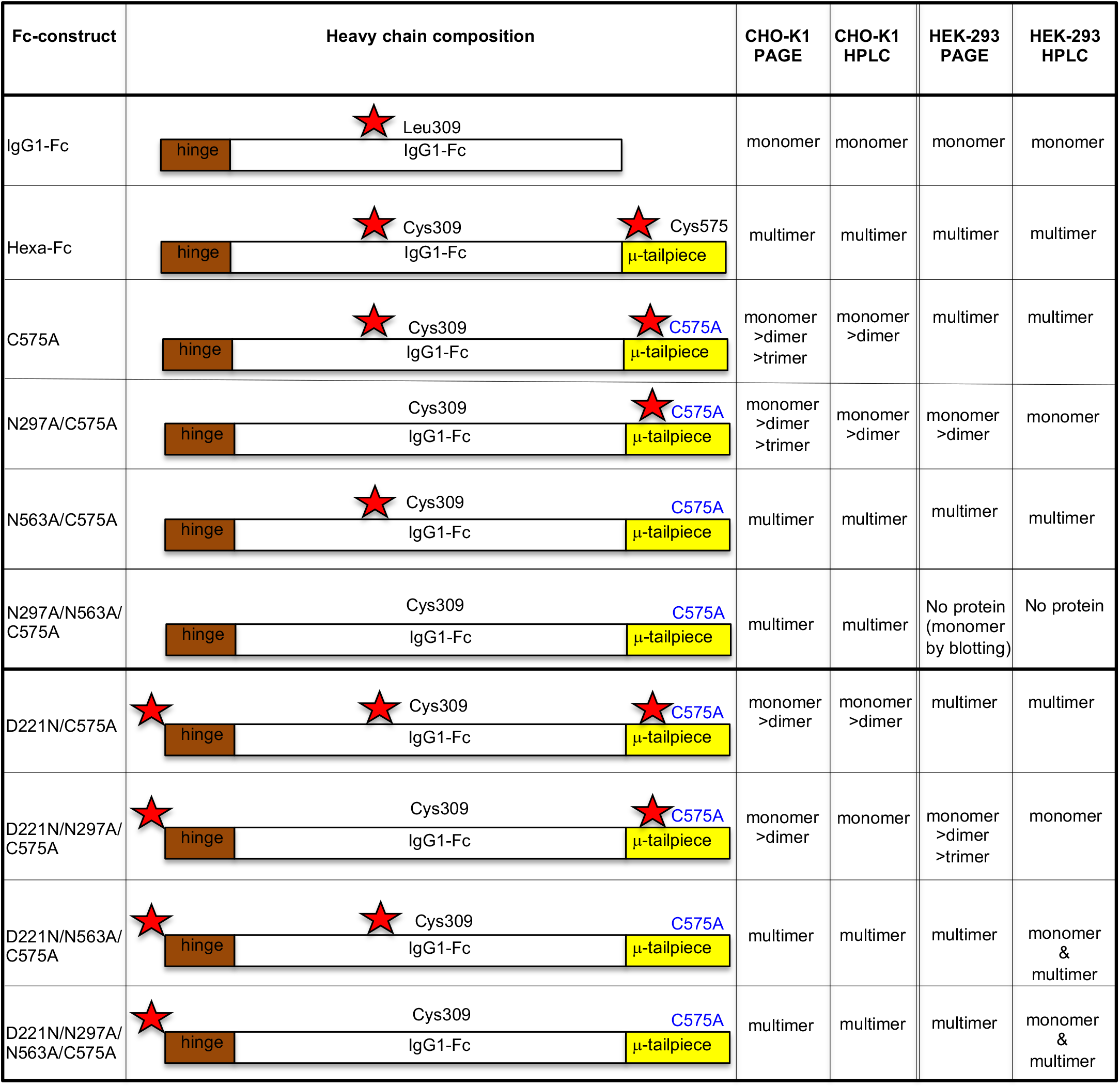
Schematic showing the various hexa-Fc glycan mutants in which Cys-575 is mutated to alanine to create the C575A panel of mutants. Stars indicate the hinge Asn-221, the Cγ2 Asn-297, and the tailpiece Asn-563 glycan sites respectively.

**FIGURE 2.**
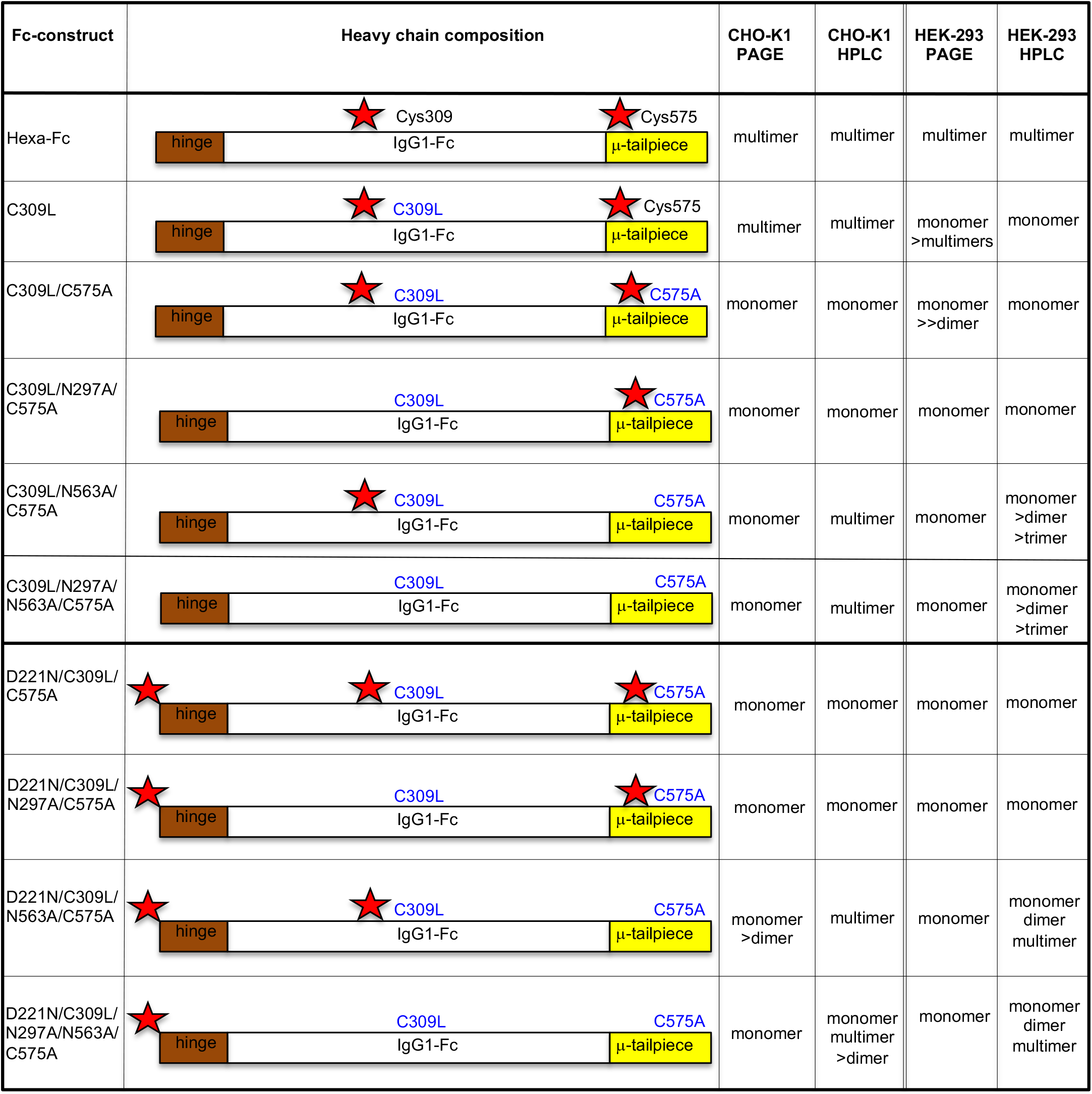
Schematic showing the C575A panel of glycan mutants from Fig. 1 in which the which Cys-309 and Leu-310 are changed to leucine and histidine, as found in the native IgG1 Fc sequence to create the C309L/C575A panel of mutants. Stars indicate the hinge Asn-221, the Cγ2 Asn-297, and the tailpiece Asn-563 glycan sites.

### Size analysis using SE-HPLC

A SEC3000 [300 × 7.8 mm] column (Beckman) was set up on a Dionex ICS3000 HPLC system and pre-equilibrated with 0.2 μm filtered PBS. Protein samples at concentrations ranging from 0.5-1 mg/mL were placed in a pre-cooled auto-sampler at 4°C and 10 μL of each was sequentially injected onto the column. Each sample was run for 1.5 column volumes in PBS at a flow rate of 0.25 mL/min. Elution was monitored at 280 and 214 nm. The column was calibrated by running standard proteins (BioRad: thyroglobulin, bovine IgG, ovalbumin, myoglobin and cyanocobalamin) under the same conditions.

### Receptor and complement binding assays

Methods describing the binding of mutants to tetrameric human DC-SIGN (Elicityl), Siglec-1, Siglec-4, and Siglec-3 (Stratech Scientific) have all been described previously (30, 31). The same ELISA protocol was used for Siglec-2, CD23, dec-1, dec-2, clec-4a, clec-4d, MBL and MMR (Stratech Scientific or Bio-Techne). Binding of C1q has been described previously (30, 31). ELISAs were used to investigate binding of Fc glycan mutants to human FcγRI, FcγRIIA, FcγRIIB, FcγRIIIA, and FcγRIIIB (Bio-Techne). Receptors were coated down onto ELISA plates (Nunc) in carbonate buffer pH 9 (Sigma-Aldrich) at 2 μg/ml overnight at 4°C, unless otherwise specified. The plates were blocked in PBS / 0.1% Tween-20 (PBST) containing 5% dried skimmed milk. Plates were washed three times in PBST before adding Fc mutant proteins at the indicated concentrations and left at 4°C overnight. Plates were washed as above and incubated for 2h with 1:500 dilution of an alkaline phosphatase-conjugated goat F(ab′)_2_ anti-human IgG (Jackson Laboratories). Plates were washed and developed for 15 min with 100 μl/well of a Sigmafast *p-*nitrophenyl phosphate solution (Sigma-Aldrich). Plates were read at 405nm, and data plotted with GraphPad Prism.

### Hemagglutination inhibition assay (HIA)

Native influenza B Hong-Kong 5/72 was obtained from Meridian Life Sciences. To determine the optimal virus-to-erythrocyte ratio, two-fold virus stock dilutions were prepared in U-shaped 96-well plates (Thermo Scientific). The same volume of a 1% human O+ red blood cell suspension (Innovative Research) was added to each well and incubated at room temperature for 1h until erythrocyte pellets had formed. After quantifying the optimal virus-to-erythrocyte concentration (4HA units), serial two-fold dilutions of Fc, control IVIG (GammaGard, Baxter Healthcare) and polyclonal goat anti-influenza B (Biorad) were prepared, all starting at a concentration of 2 μM, and mixed with 50 μl of the optimal virus dilution. After 30 min incubation at room temperature, 50 μl of the human erythrocyte suspension was added to all wells, and plates incubated at room temperature for 1h, after which erythrocyte pellets could be observed in the positive controls and positive samples.

### Binding to FcγRs by Biacore

Binding to FcγRs was carried out using a Biacore T200 biosensor (GE Healthcare). Recombinantly expressed FcγRS (R&D systems or Sino Biologicals) were captured via their histidine tags onto CM5 chips pre-coupled with ~9000 reflective units anti-His Ab (GE Healthcare) using standard amine chemistry. Fc mutants were injected over captured receptors at a flow rate of 20 μl/min, and association and dissociation monitored over indicated time scales before regeneration with two injections of glycine (pH 1.5) and recalibration of the sensor surface with running buffer (10 mM HEPES, 150 mM NaCl [pH 7]). Assays were visualized with Biacore T200 evaluation software v2.0.1.

### N-glycomic analysis

N-glycomic analysis was based on previous developed protocol with some modifications (32). Briefly, the N-glycans from 50 μg of each sample were released by incubation with NEB Rapid™ PNGase F and isolated from peptides using Sep-Pak C18 cartridges (Waters). The released N-glycans were permethylated, prior to Matrix-assisted laser desorption ionization (MALDI) MS analysis. Data were acquired using a 4800 MALDI-TOF/TOF mass spectrometer (Applied Biosystems) in the positive ion mode. The data were analyzed using Data Explorer (Applied Biosystems) and Glycoworkbench (33). The proposed assignments for the selected peaks were based on composition together with knowledge of biosynthetic pathways.

## Results

### Disulfide bonding and glycosylation influence the multimerization states of Fc mutants expressed by HEK 293-F cells

Two panels of glycosylation- and cysteine-deficient mutants previously expressed by CHO-K1 cells were generated in HEK 293-F cells (Figs. 1 and 2). As observed with CHO-K1 cells, the HEK 293-F cells were capable of making all the mutants to high yields (~30 mg/L) with the exception of the N297A/N563A/C575A mutant for which we were unable to generate sufficient protein for further work. Generally, all the mutants migrated on SDS-PAGE with the expected molecular weights for their glycosylation or disulfide bonding states (Fig. 3), as previously described for the same mutants expressed by CHO-K1 cells (23).

**FIGURE 3.**
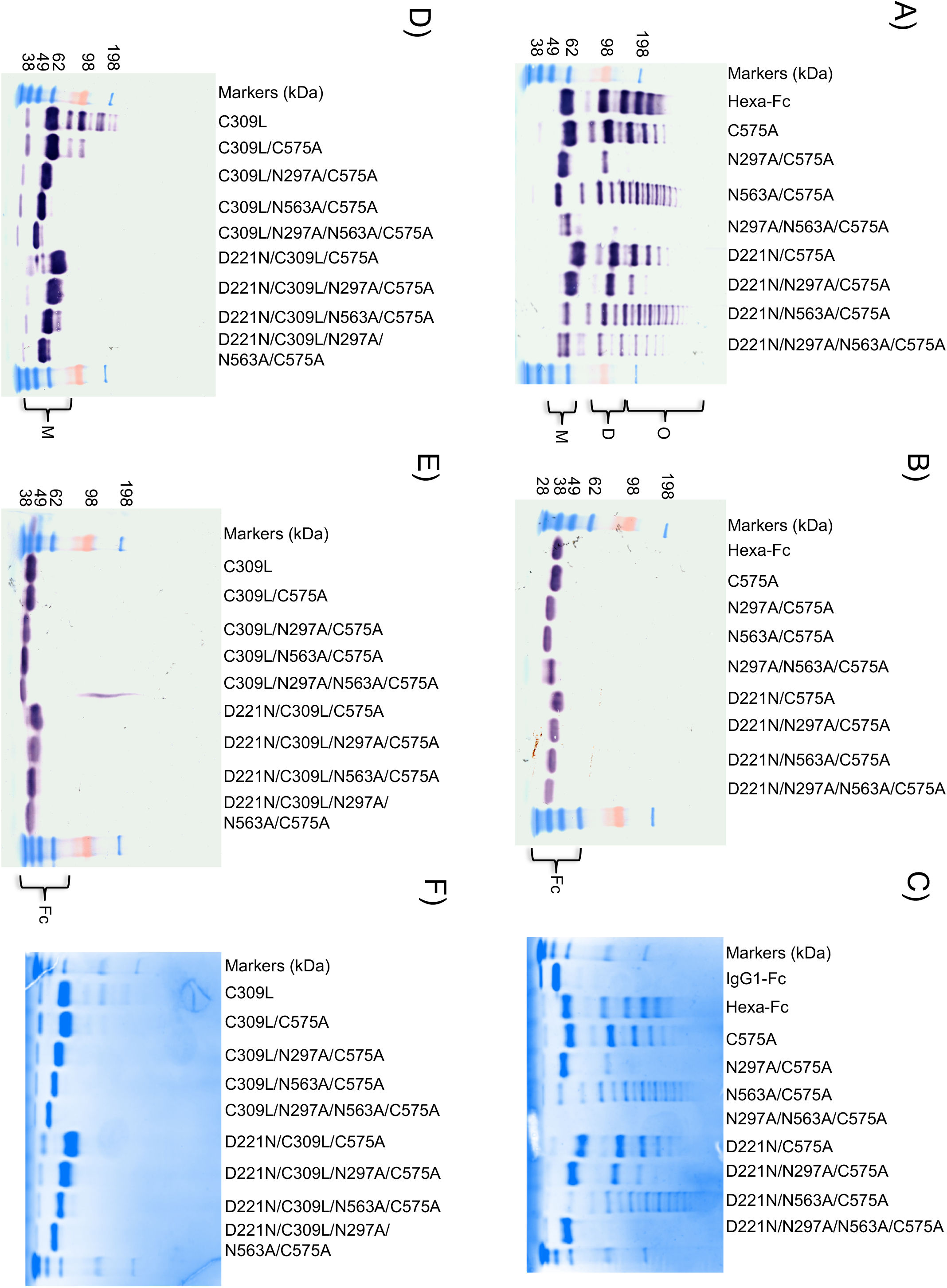
Characterization of mutant Fc proteins by SDS-PAGE. (**A**) Cys-309 competent mutants in which Cys-575 is mutated to alanine to create the C575A panel of mutants. Mutants with N563A run as laddered multimers. Insufficient material was obtained with N297A/N563A/C575A for further analysis. The addition of the N-X*-*(T/S) glycan sequon to generate N-terminally glycosylated hinges (the D221N series of mutants) did not affect multimerization but rather increased the molecular mass of all mutants. The N297A mutants run as monomers, dimers and trimers. (**B**) the same mutants as in (A) but run under reducing conditions. The D221N/C575A mutant has the largest mass because it has three glycans attached. The types of glycans attached at Asn-221, Asn-297, and Asn-563 for all the mutants are shown in Fig. 9 and supplemental figures. The decreasing molecular masses seen in the Fc represent the sequential loss of N-linked glycans. (**C**) The same mutants as in (A) but stained with Coomassie reagent. (**D**) Substitution of Cys-309 with leucine onto the C575A mutants shown in (A) to create the double cysteine knockouts, which all run as monomers. C309L in which Cys-575 is present also multimerizes. (**E**) The same mutants as in (D) but run under reducing conditions. Note that the D221N/C309L/C575A mutant with three glycan sites has the largest mass, as seen with the equivalent mutant D221N/C575A in panel (A). (**F**) Coomassie-stained gel of (D). All proteins were run under either non-reducing (panels A and D) or reducing conditions (panels B and E) at 2 μg protein per lane on 4-8% acrylamide gradient gels, transferred to nitrocellulose, and blotted with anti-human IgG Fc (Sigma-Aldrich).

In an earlier study with CHO-K1 cells we demonstrated that a proportion of molecules in which the tailpiece Asn-563 glycan was substituted for alanine ran as multimers in solution when examined by SE-HPLC (24). The loss of the bulky Asn-563 glycan exposes hydrophobic amino acid residues in the tailpiece that facilitate non-covalent interactions in solution. Such N563A-dependent multimerization was also observed with mutants expressed by HEK 293-F cells, although the proportion of multimers to monomers (with the notable exception of the N563A/C575A mutant) was generally lower when mutants were made by this cell line (Fig. 4). Clearly, the choice of cell line and consequently the types of post-translational modifications, dramatically impact on the biophysical properties of these molecules in solution.

**FIGURE 4.**
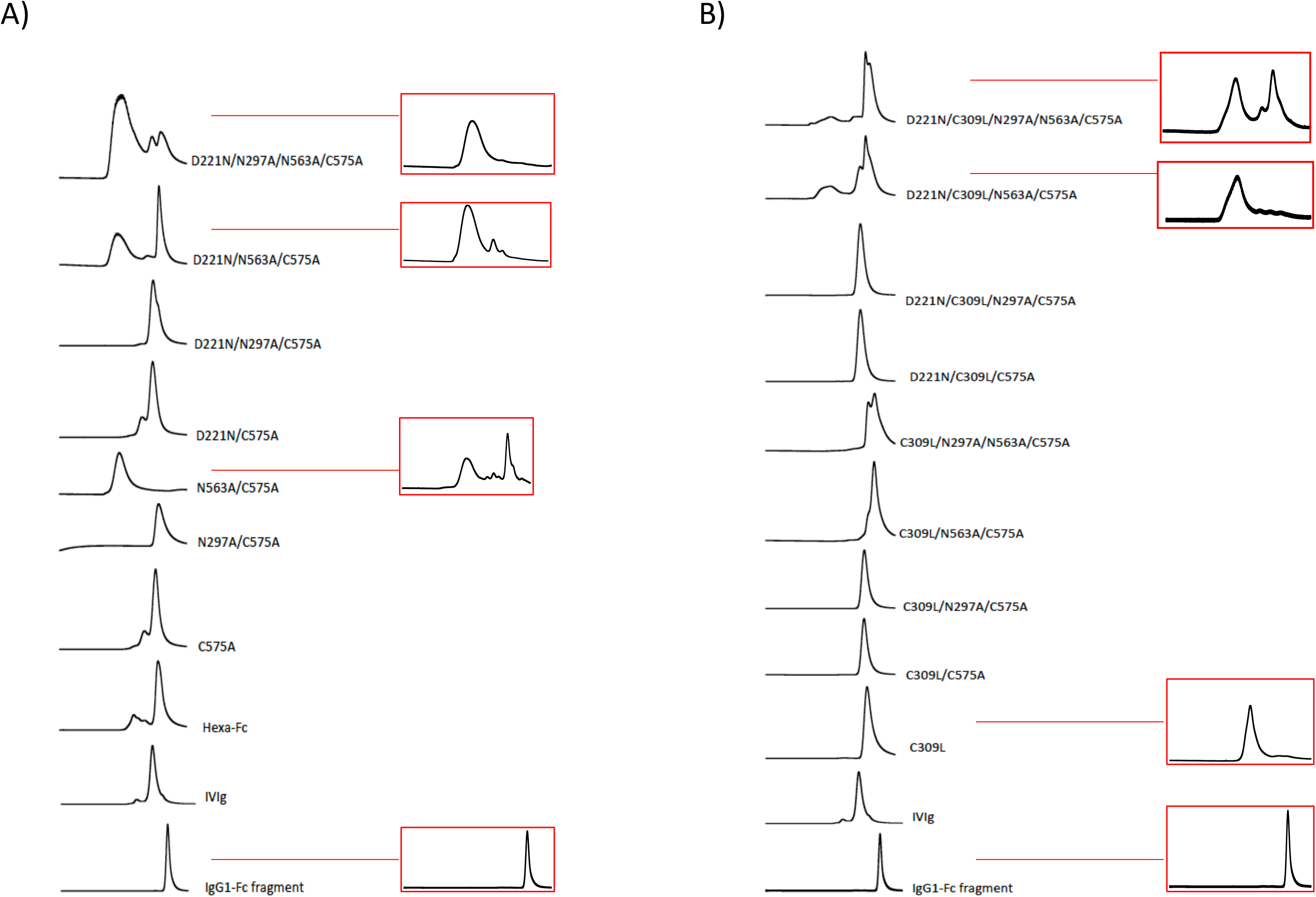
Size-exclusion chromatography analysis of Fc mutants expressed in HEK 293-F cells. (A) The C575A panel of mutants. (B) The C309L/C575A panel of mutants. Boxed chromatograms represent profiles for equivalent mutants expressed in CHO-K1 cells, as published previously (24).

### Fc glycan mutants expressed by HEK 293-F cells show important differences in binding to glycan receptors when compared to CHO-K1 proteins

To determine the impact of the cell line on receptor binding by the two panels of Fc mutants, we investigated their interaction with soluble recombinant glycan receptors by ELISA (Fig. 5). For most of the Fc mutants, including hexa-Fc, C575A, N297A/C575A, D221N/N297A/N563A/C575A, C309L/C575A, D221N/C309L/C575A, D221N/C309L/N297A/C575A and D221N/C309L/N297A/N563A/C575A, expression in HEK 293-F cells reduced the binding to glycan receptors when compared to equivalent molecules expressed in CHO-K1 cells (Fig. 5). However, two Fc mutants (D221N/N563A/C575A and D221N/C309L//N573A/C575A) were notable for their enhanced binding to all the glycan receptors investigated when expressed in HEK 293-F cells. Given that both mutants multimerize poorly by comparison to the equivalent mutants made in CHO-K1 cells (Fig. 4), we attribute this enhanced glycan receptor binding not to increased avidity effects but to fine differences in the attached glycan structures. Therefore, the choice of cell line can dramatically impact on the ability of individual Fc mutants to interact with glycan receptors.

**FIGURE 5.**
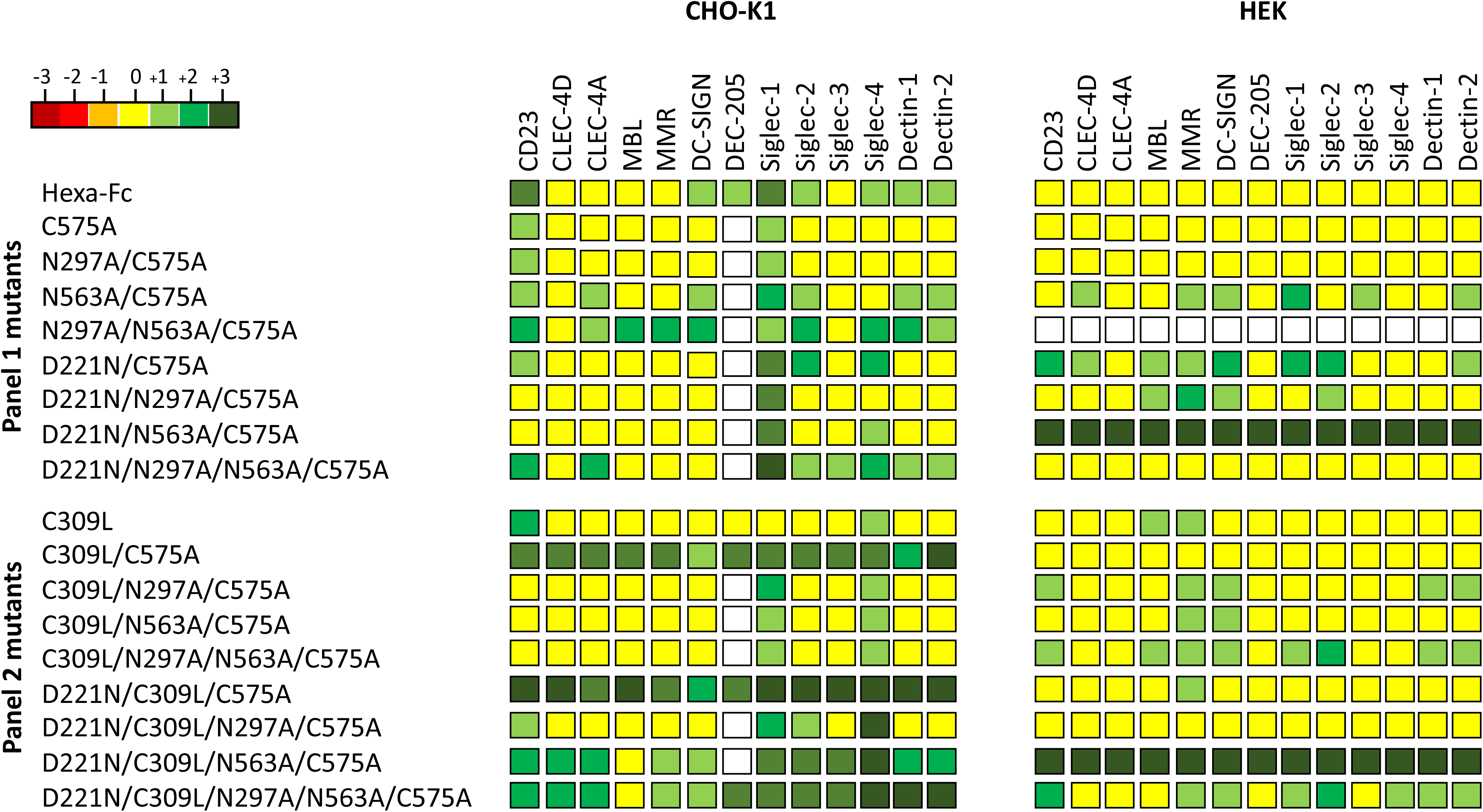
Shading matrix showing the differential binding of HEK 293-F or CHO-K1 mutant proteins to recombinant glycan receptors. Results from at least two independent ELISA experiments are expressed as fold change (up or down) with respect to the internal IgG1 Fc control run on each plate. Standalone ELISA data are provided in the supplementary figures. White boxes = not tested.

### Fc glycan mutants expressed by HEK 293-F cells show important differences in binding Fcγ-receptors compared to CHO-K1 cell proteins

Given the observed differences in binding to glycan receptors of the same Fc mutants expressed by two different cell lines, we also tested the impact of cell line on binding to the classical human FcγRs (Fig. 6).

**FIGURE 6.**
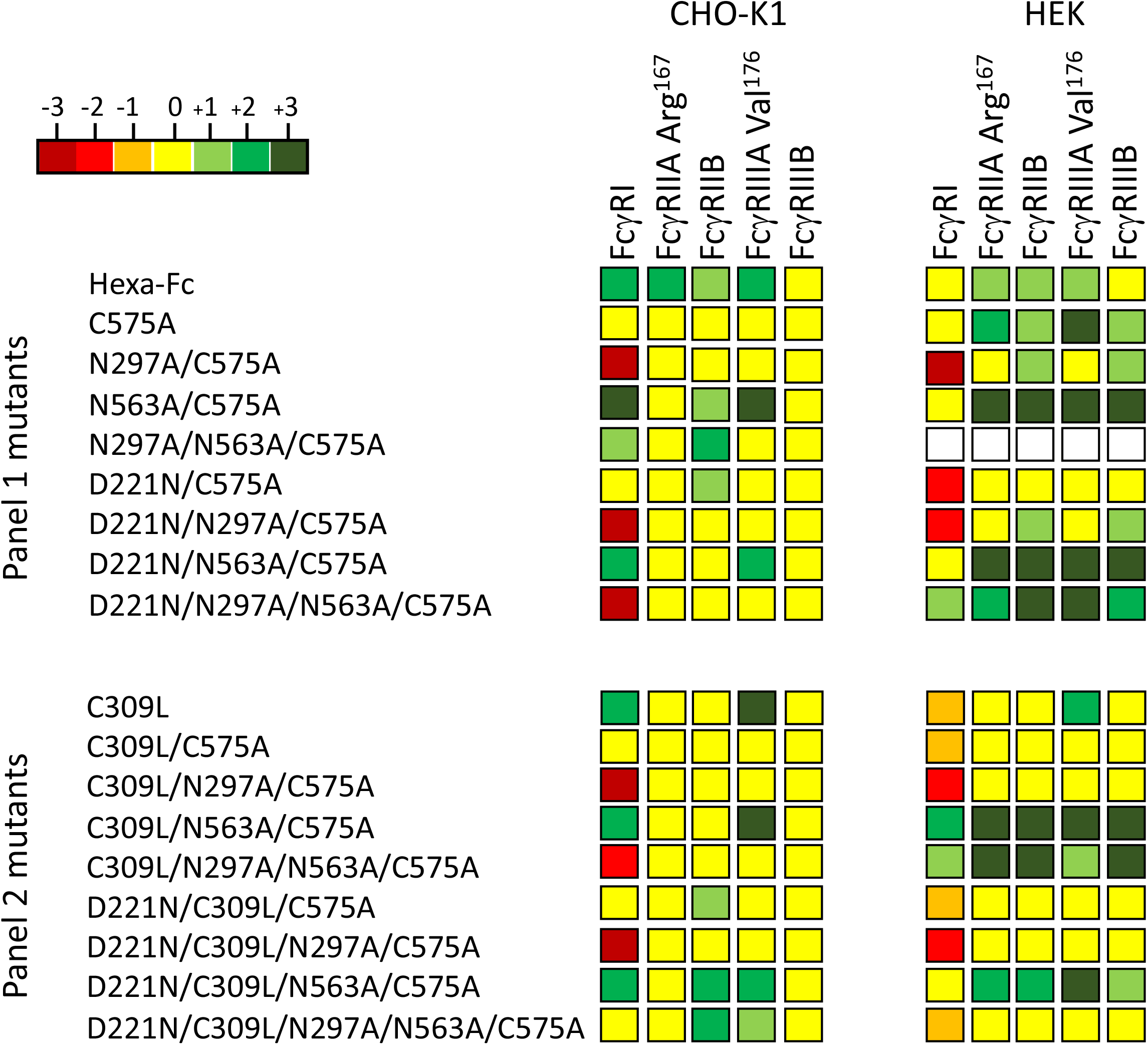
Shading matrix showing the differential binding of HEK 293-F or CHO-K1 mutant proteins to recombinant Fcγ receptors. Results from at least two independent ELISA experiments are expressed as fold change (up or down) with respect to the internal IgG1 Fc control run on each plate. Standalone ELISA data are provided in the supplementary figures.

The most significant difference observed was the ability of certain HEK expressed mutants (N563A/C575A, D221N/N563A/C575A, D221N/N297A/N563A/C575A, C309L/N563A/C575A, C309L/N297A/N563A/C575A and D221N/C309L/N563A/C575A) to bind human FcγRIIA (Arg-167) and FcγRIIIB. This is in stark contrast to the same proteins expressed in CHO-K1 cells, where not one single mutant from each panel bound either of the two low-affinity receptors (Fig. 6 and (24)).

To examine the interaction with human FcγRIIA (Arg-167) and FcγRIIIB in more detail, we tested binding of two of these mutants (C309L/N563A/C575A and D221N/C309L/N563A/C575A) to FcγRIIA (Arg-167) and FcγRIIIB by surface plasmon resonance analysis (Fig. 7). Both mutants displayed slower apparent off rates compared to the control IgG1-Fc monomer, consistent with avidity effects either through binding to multiple immobilized FcγR molecules or rebinding effects (Fig. 7). Therefore, the choice of cell line impacts on the ability of individual Fc mutants to interact with FcγRs, and in particular FcγRIIA (Arg-167) and FcγRIIIB.

**FIGURE 7.**
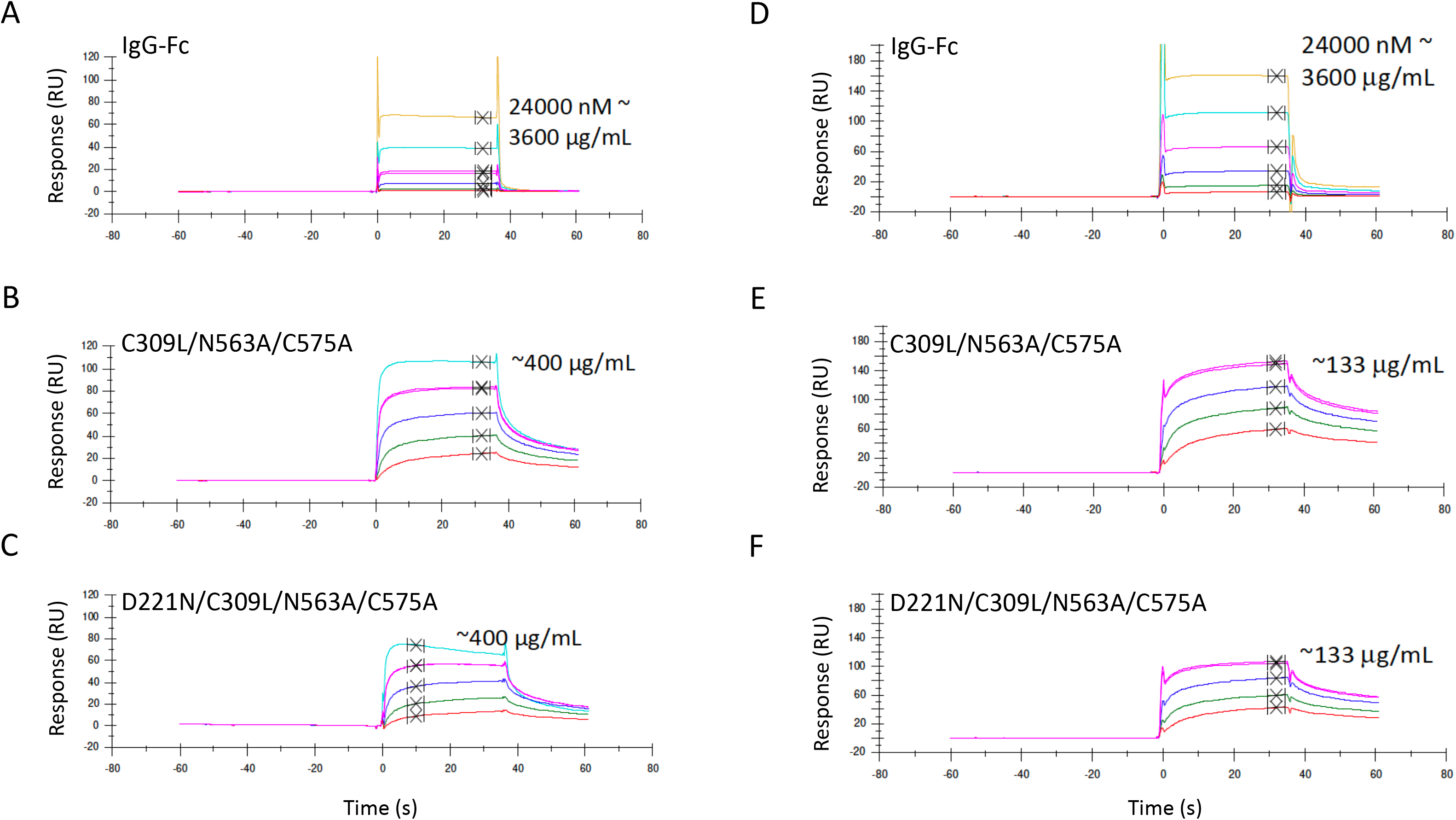
Surface plasmon resonance analysis. Binding of selected mutants to human FcγRIIIB (panels **A-C**) or FcγRIIA-Arg^167^ (panels **D-F**) by Biacore. Control IgG1 Fc (panels **A,D**) is compared to C309L/N563A/C575A (panels **B,E**) and D221N/C309L/N563A/C575A (panels **C,F**). Curves show doubling dilutions from the highest indicated concentration of protein. Because of the varying stoichiometry of the molecules shown (as seen in Figs. 3 and 4), an accurate determination of the interaction kinetics is not possible. Binding was to receptors sourced from R&D systems (Bio-Techne).

### Fc glycan mutants expressed by HEK 293-F cells show improved binding to human C1q

An important functional and safety attribute for therapeutic administration of Fc fragments is their ability to bind C1q, and thus initiate the classical pathway of complement activation. Binding of C1q was assessed by ELISA to selected mutants expressed from each cell line (Fig. 8). Mutants D221N/C575A, D221N/C309L/C575A, C309L/N297A/N563A/C575A and C309L/N563A/C575 expressed in HEK 293-F cells showed improved binding to C1q, compared to their counterparts expressed in CHO-K1 cells, and no change in binding in either direction was observed for IgG1-Fc, D221N/N563A/C575A, D221N/N297A/C575A, D221N/C309L/N297A/C575A, D221N/C309L/N297A/N563A/C575A, and C309L/N297A/C575A (Fig. 8).

**FIGURE 8.**
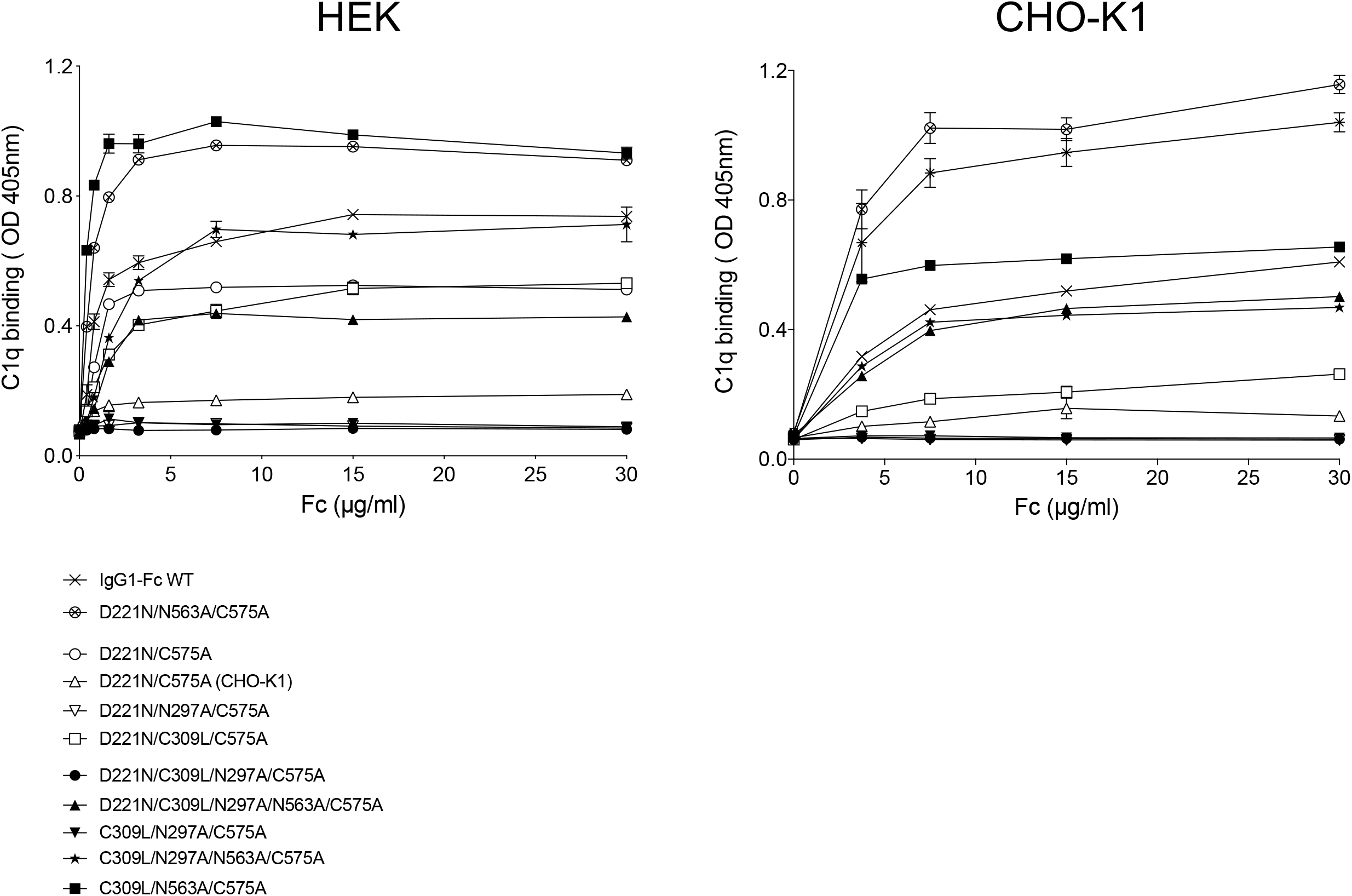
Binding of selected C575A and C309L/C575A mutants to complement component C1q. Mutants expressed in HEK 293-F cells bind human C1q better than the equivalent mutants expressed in CHO-K1 cells. Compare for example the D221N/C575 mutant made in CHO-K1 cells (open triangle) against the same mutant made in HEK 293-F cells (open circle) and compared on the same plate. Error bars represent standard deviations around the mean value, n=2 independent ELISA experiments.

Both the D221N/C575A and D221N/C309L/N297A/C575A mutants from CHO-K1 cells have been shown previously to block influenza-mediated hemagglutination (ref 23, and Fig. 11 below), and thus D221N/C575A expressed in HEK 293-F cells that binds C1q may not be favored for clinical development over the same molecule expressed by CHO-K1 cells (27).

### Fc glycan mutants expressed in HEK 293-F cells have more complex glycosylation profiles than the equivalent mutants expressed in CHO-K1 cells

The structure of the N-glycan on the Fc of IgG antibodies has been shown to influence multiple receptor interactions (3, 34, 35). Unlike the relatively simple glycosylation of the Fc mutants previously described for CHO cells (23, 24), HEK cells are capable of producing more complex N-glycan structures on their glycoproteins (36).

We investigated the nature of the N-glycans on the two panels of glycosylation- and cysteine-deficient mutants by MALDI-TOF mass spectrometry-based glycomic analysis (complete data set for both panels of mutants provided in supplementary figures). A core-fucosylated biantennary structure without antennary galactosylation, *m/z* 1835 (GlcNAc_4_Man_3_Fuc_1_), is the base peak of spectra from all IgG1-Fc mutants produced by HEK cells (Fig. 9 and supplementary figures).

**FIGURE 9.**
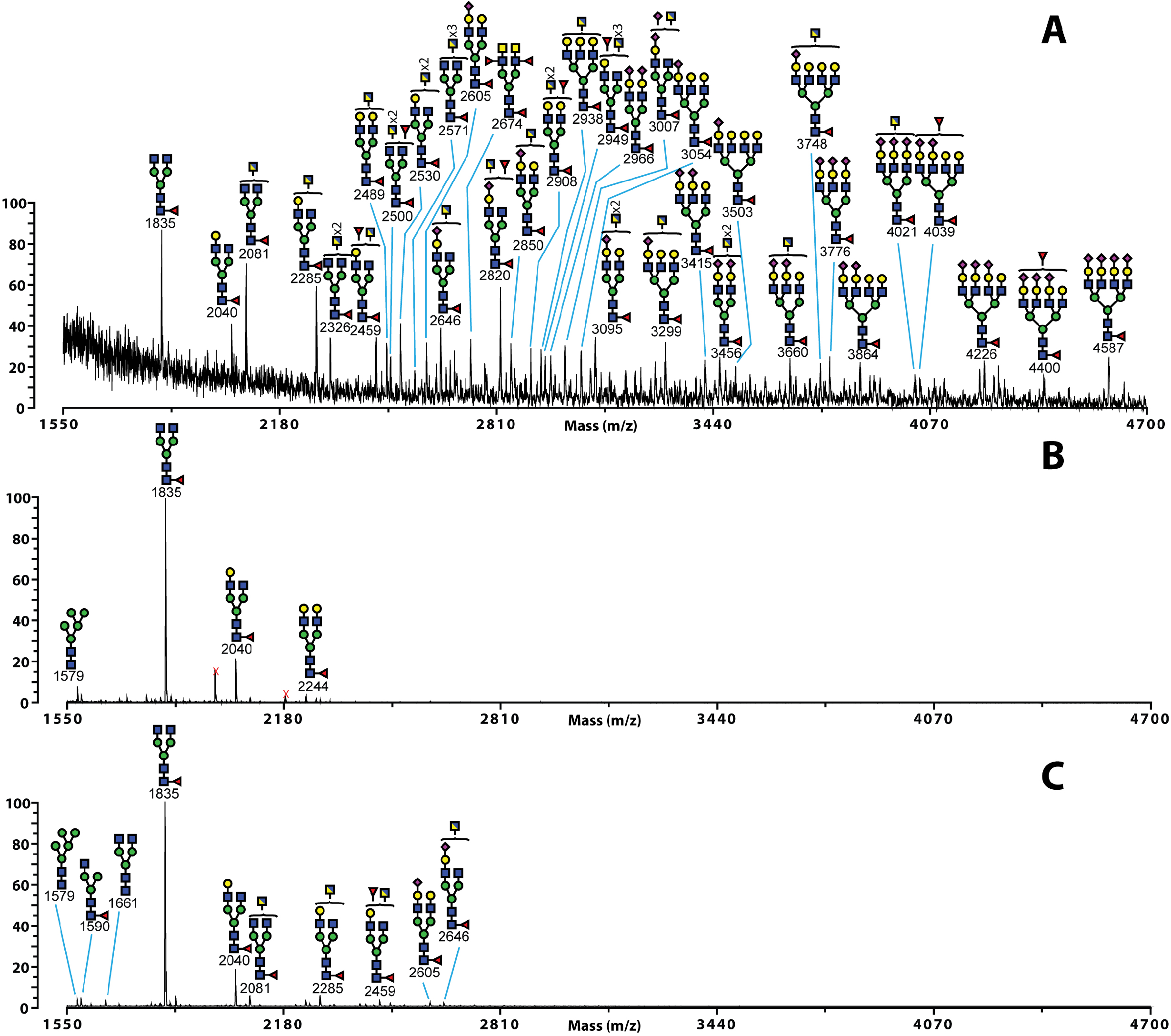
MALDI-TOF MS profiles of permethylated N-glycans from the N297A/C575A (**A**), N563A/C575A (**B**), and D221N/N297A/N563A/C575A (**C**) Fc glycan mutants. Linkage determined monosaccharides are positioned above the bracket on a structure. Poly-hexose contaminants are highlighted with crosses. The data were acquired in the positive ion mode to observe [M^+^Na]^+^ molecular ions. All the structures are based on composition and knowledge of N-glycan biosynthetic pathways. Structures shown outside a bracket have not had their antenna location unequivocally defined.

The types of site-specific glycans attached to either Asn-221, Asn-297 or Asn-563 could be determined using both the C575A or C309L/C575A panels of mutants. For example, only sugars attached to Asn-297 are available for sampling in either the N563A/C575A or C309L/N563A/C575A, mutants that therefore also allow the contribution of disulfide bonding to glycosylation at Asn-297 to be elucidated.

The N-glycosylation of Asn-297 is dominated by core-fucosylated bi-antennary glycans (*m/z* 1835 and 2040) with varied galactosylation levels (Gal_0-2_GlcNAc_4_Man_3_Fuc_1_), and a Man_5_GlcNAc_2_ (*m/z* 1579) high mannose structure is also observed (Fig. 9B). The Asn-563 N-glycans are much more complex and heterogeneous. Abundant truncated structures at m/z 2081 and 2285 have potentially terminal GlcNAc or GalNAc (Fig. 9A). Antennal fucosylation and sialylation is also observed on structures which can assemble sialyl lacNAc, sialyl-Lewis x/a, fucosylated LacdiNAc or sialylated LacdiNAc (GalNAc-GlcNAc), for example peak *m/z* 4039 (NeuAc_2_Gal_4_GlcNAc_6_Man_3_Fuc_2_). The presence of *m/z* 2674 (GalNAc_2_GlcNAc_4_Man_3_Fuc_3_), in the N297A/C575A mutant confirms the presence of fucosylated LacdiNAc epitopes on the Asn-563 site. Thus, glycosylation at Asn-563 is different to that seen from CHO-K1 cells that assemble less diverse structures without antennal fucosylation and therefore more terminal sialyl-LacNAc (23, 24).

The Asn-221 site is mainly composed of bi-antennary complex structures (Fig. 9C). Excluding the base peak, four structures in the C575A background (m/z 2081, 2285, 2459 and 2646) or five structures in the C309L/C575A background (2081, 2285, 2459, 2489 and 2734) could form LacdiNAc antenna (GalNAc-GlcNAc). Antennal fucosylation and sialylation is also observed (Fig. 9C and supplementary Figures).

In summary these data show that the types of glycans attached to either Asn-221, Asn-297 or Asn-563 are different between cell lines but are not grossly affected by disulfide bonding.

### Fc glycan mutants expressed in HEK 293-F cells are less sialylated than the equivalent mutants expressed in CHO-K1 cells

Site-specific levels of sialylation were semi-quantitatively assessed for both panels of mutants and compared to levels seen in the equivalent mutants expressed in CHO-K1 cells (Fig. 10). Although levels of sialylated glycans attached at positions Asn-297 (the N563A/C575A mutant) and Asn-563 (the N297A/C575A mutant) are similar for both cell lines (Fig. 10), a marked reduction in levels of sialylated glycans at Asn-221 (the D221N/N297A/N563A/C575A mutant) is observed when this mutant is expressed in HEK cells (2.8% against 81.8% in CHO, Fig. 10). Removal of Asn-297 generally enhanced levels of sialylation at both Asn-221 and Asn-563, irrespective of the cell line or the multimerization state of the proteins, e.g. compare N297A/C575A vs. C575A and C309L/N297A/C575A vs. C309L/C575A (Fig. 10). The choice of cell line therefore dramatically affects the overall levels of sialylation at individual N-linked attachment sites within the glycan-modified Fc variants.

**FIGURE 10.**
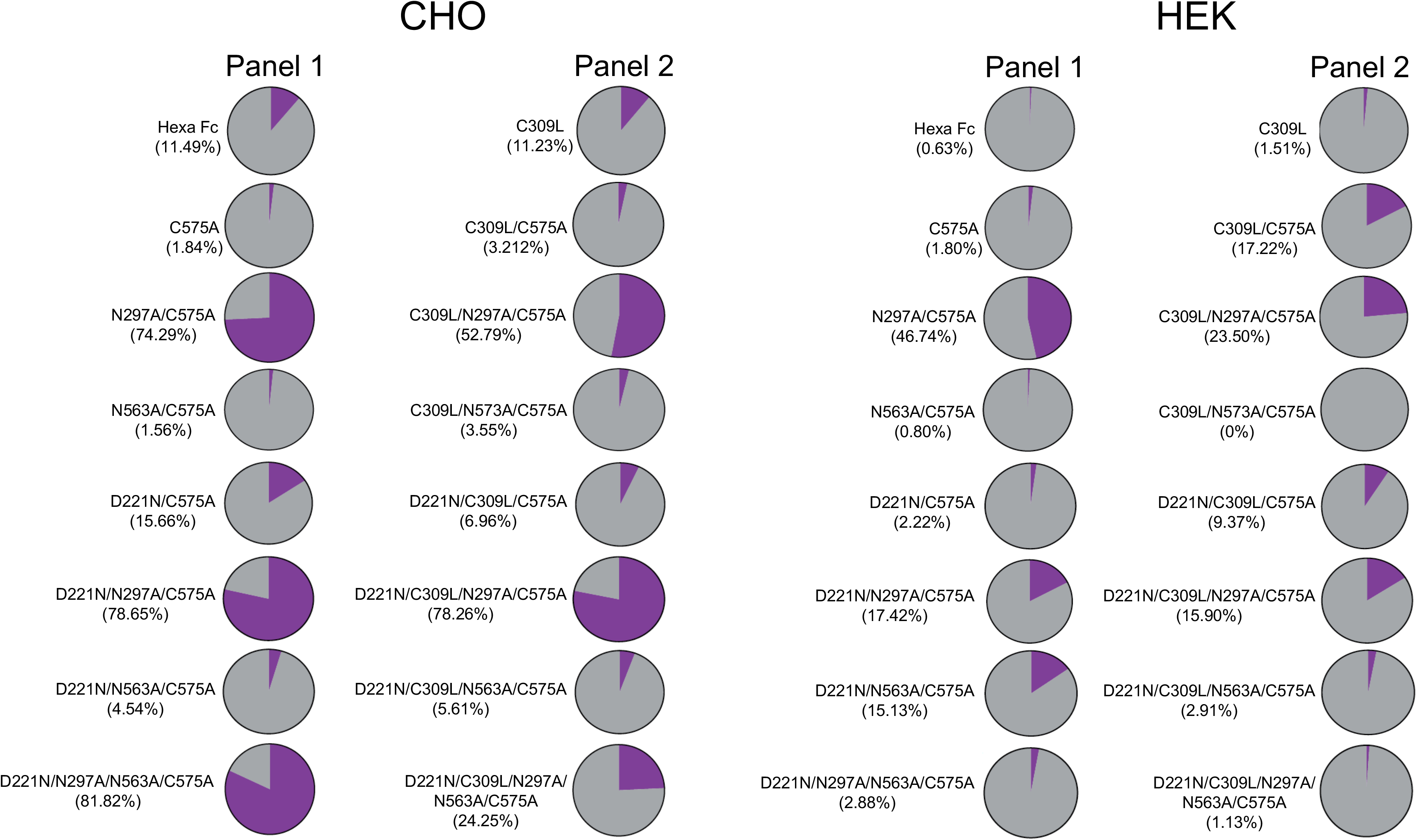
Semi-quantitative determination of sialylated (purple) against neutral (grey) glycans from the C575A and C309L/C575A mutants expressed in CHO-K1 or HEK 293-F cells. Values shown in brackets under the names of each mutant show percentage sialylated structures as determined from summed intensities.

**FIGURE 11.**
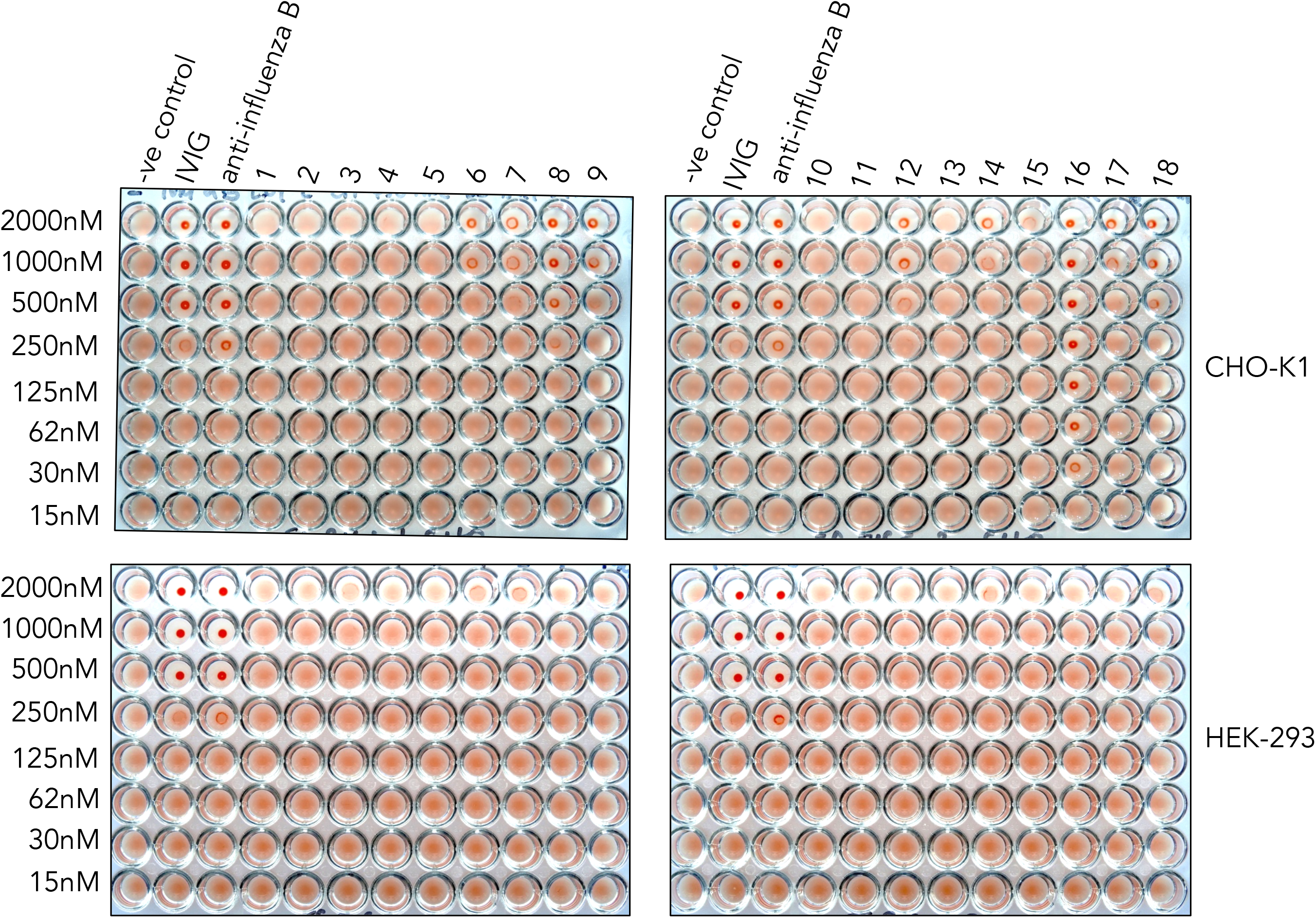
Impact of Fc glycosylation on influenza B-mediated hemagglutination. Mutant Fcs manufactured in either HEK 293-F or CHO-K1 cells were compared to equimolar concentration of IVIG or polyclonal anti-influenza B antibodies at inhibiting virus-mediated agglutination of human erythrocytes. 1. L309C, 2. C575A, 3. N297A/C575A, 4. N563A/C575A, 5. No protein, 6. D221N/C575A, 7. D221N/N297A/C575A, 8. D221N/N563A/C575A, 9. D221N/N297A/N563A/C575A, 10. C309L, 11. C309L/C575A, 12. C309L/N297A/C575A, 13. C309L/N575A/C575A, 14. C309L/N297A/N563A/C575A, 15. D221N/C309L/C575A, 16. D221N/C309L/N297A/C575A, 17. D221N/C309L/N563A/C575A, 18. D221N/C309L/N297A/N563A/C575A. A constant amount of influenza B Hong-Kong 5/72 virus was incubated with titrated amounts of the Fc glycan mutants and added to human O+ erythrocytes that were then allowed to sediment at room temperature for 1h. Non-agglutinated RBCs form a small halo. n=3 independent experiments.

### Asn-221-containing mutants are poor inhibitors of hemagglutination by influenza virus when expressed in HEK 293-F cells

To test if the choice of cell line affected the functionality of the two panels of mutant Fcs, we used the World Health Organization (WHO) hemagglutination inhibition assay (HIA) to quantify the inhibitory titers for each mutant against an influenza B virus (Fig. 11). As shown previously with an avian influenza A (H1N1) (24), mutants containing Asn-221 hinge-attached glycans, and in particular the D221N/C309L/N297A/C575A mutant, prevented hemagglutination by an influenza B virus at concentrations as low as 30nM, an eight-fold improvement over equimolar IVIG or polyclonal anti-influenza B antisera (Fig. 11). In stark contrast, the same mutants expressed by HEK 293-F cells were unable to inhibit hemagglutination by either influenza A (not shown) or influenza B virus (Fig. 11). This shows that the functional potential of individual glycan-modified Fc mutants is dependent on the choice of cell line used for their manufacture.

## Discussion

We have shown using CHO-K1 cells that the structure and effector function of human IgG1-Fc can be profoundly altered by the addition or removal of N-linked glycosylation (23, 24). For example, we could show that Fc fragments containing complex biantennary glycans attached to both the N- and C-terminal ends of the Fc could inhibit influenza A-mediated agglutination of human erythrocytes (24). The aim of the current study was to reveal possible variation in functional glycosylation related to differences in two host cell lines, CHO-K1 and HEK 293-F, particularly as antibodies and Fc fusions are the fastest growing therapeutic class in the pharmaceutical industry (26, 37, 38).

Two intriguing aspects of N-linked glycosylation are relevant to this study. First, the differential binding seen to human glycan (Fig. 5) and Fcγ (Fig. 6) receptors between the same mutants expressed by two different cell lines. These differentially manufactured mutants now need to be compared in relevant *in vivo* disease models where the Fc is therapeutically useful, given that differential sialic acid linkages, α2,6 and α2,3, are known to impact on the anti-inflammatory properties of the Fc (39, 40). Such nuanced glycosylation may also explain why the therapeutic efficacy of molecules generated by different expression systems, and subsequently tested in different animal models, do not always translate to efficacy in human studies (41).

Second, we have studied the exquisite impact of the host cell line on the efficacy of sialylated Fcs to inhibit influenza viruses (Fig. 11). One possible explanation is that overall sialylation levels for all the influenza blocking mutants, in particular the D221N/C309L/N297A/C575A mutant, are approximately five-fold lower when expressed by HEK 293-F cells (Fig. 10). However, overall level of sialylation is not the only possible explanation for the relative efficacy of the CHO-K1 mutants in inhibiting influenza virus hemagglutination, as the CHO-K1-expressed D221N/C575A mutant also contained approximately five-fold less sialyation than the D221N/C309L/N297A/C575A mutant made in the same cell line (Fig. 10). This indicates that the fine specificity (e.g. α2,3 vs. α2,6 linkages) of these sialylated glycans may also be a contributing factor to their efficacy.

As demonstrated previously for influenza A (24), binding and inhibition of influenza B viruses is stronger with mutants containing Asn-221, and in particular by the monomeric mutant D221N/C309L/N297A/C575A in which the N- and C-terminal sialylated sugars are spaced ~60Å apart (Figs. 11 and (24)). Recent biophysical studies with alternative glycan-decorated scaffolds have shown that ~1,000 fold enhancements over monovalent binding to HA can be achieved with only two sialylated ligands, provided the sugars are arranged 50-100Å apart (42, 43). As we also observed with Fc multimerizing mutants from panel 1, no additional benefit with respect to virus neutralization was gained with larger, more complex sialylated structures (Fig. 11, and as seen with the D221N/N297A/N563A/C575A mutant).

We do not yet know if sialylated Fcs are susceptible to cleavage by the viral neuraminidase. Although a decoy for NA may be a therapeutically attractive strategy (44), we have not observed a direct decay in the HIA after prolonged incubation. This suggests that the high specific avidity of these molecules for HA may reduce their susceptibility to NA, a hypothesis that fits with the relatively low efficiency of neuraminidase (*k*_cat_ = 30-155^**s-1**^), together with the asymmetric distribution of NA in relation to HA on the surface of filamentous influenza viruses (45–47).

In order to be useful compounds when administered intranasally, or as an aerosol, the sialylated Fc needs to out-compete sialylated mucins that viruses use through ratchet-like interactions with HA and NA to migrate to the underlying respiratory epithelium (45). Of the 15 known human mucins in the human lung, only MUC5 has been shown to protect from influenza (48, 49). Most sialic acid found on human mucins are O-glycosylated, and where N-linked attachments do occur, these are mostly sialylated via α2,6-linkages (49). Thus, we were surprised that none of the Fc leads inhibited influenza A (H1N1 propagated in hen eggs) or influenza B (Hong Kong 5/72 propagated in MDCK cells) agglutination of human O^+^ erythrocytes when manufactured by HEK 293-F cells that attach more human type α2,6-linked sialic acid (Fig. 10).

The apparent importance of α2,3-linked N-glycans to inhibition of both influenza A and B by the CHO-K1 Fc mutants indicates that viruses can evolve away from inhibition by mucus whose predominant O-linked glycans are mostly α2,6-linked. Our working hypothesis is that HEK-expressed compounds may therefore inhibit influenza viruses that circulate in human populations or that are propagated in cell lines that attach more human-like α2,6-linked sialic acid.

Consequently, by careful consideration of the cell line used in their manufacture, new glycan repertoires with desirable binding attributes and functionality can be imparted to the therapeutically attractive Fc molecule.

Fc: Fragment crystallizable
IVIG: Intravenous Immunoglobulin
ITP: Idiopathic Thrombocytopenic Purpura
tp: tailpiece
Siglec: Sialic acid-binding immunoglobulin-type lectin
CD: Cluster Designation
CHO: Chinese Hamster Ovary
DC-SIGN: Dendritic Cell-Specific Intercellular Adhesion Molecule-3-grabbing Non-Integrin
DCIR: C-type Lectin Dendritic Cell Immunoreceptor
CLEC: C-type Lectin
HA: Hemagglutinin
HIA: Hemagglutination Inhibition Assay
HEK 293-F: Human Endothelial Kidney
MBL: Mannose-Binding Lectin
MMR: Macrophage Mannose Receptor
mAbs: monoclonal Antibody
MALDI: matrix-assisted laser desorption ionisation
TOF: time-of-flight
SE-HPLC: Size Exclusion-High Performance Liquid Chromatography

## Acknowledgements

We thank Abzena PLC for running the surface plasmon resonance analysis and Dr Mark Wilkinson for running SE-HPLC samples.

## Author contributions

R.J.P. conceived and designed the overall study. R.J.P, P.A.B., D.L., AD and SMH designed and performed experiments. R.J.P. wrote the manuscript, and all authors commented on drafts and reviewed the final manuscript.

## Disclosures

R.J.P. and P.A.B. declare that work discussed within is subject to ongoing patent applications.

## Notes

This work was supported by Pathfinder and Innovator grants from the Wellcome Trust (109469/Z/15/Z and 208938/Z/17/Z) and Institutional Strategic Support Fund (ISSF) 109469/Z/15/Z, 208938/Z/17/Z, 097830/Z/11/Z from the Wellcome Trust and MRC Confidence in Concept award MC_PC_12017 respectively. Also, by the Biotechnology and Biological Sciences Research Council grant BBF0083091 (A. Dell and S.M. Haslam).

