## Supplemental Figures for "Choice of host cell line is essential for the functional glycosylation of the fragment crystallizable (Fc) region of human IgG1 inhibitors of influenza B viruses"

### Glycan receptor binding data for series 1 mutants – methods described as per manuscript

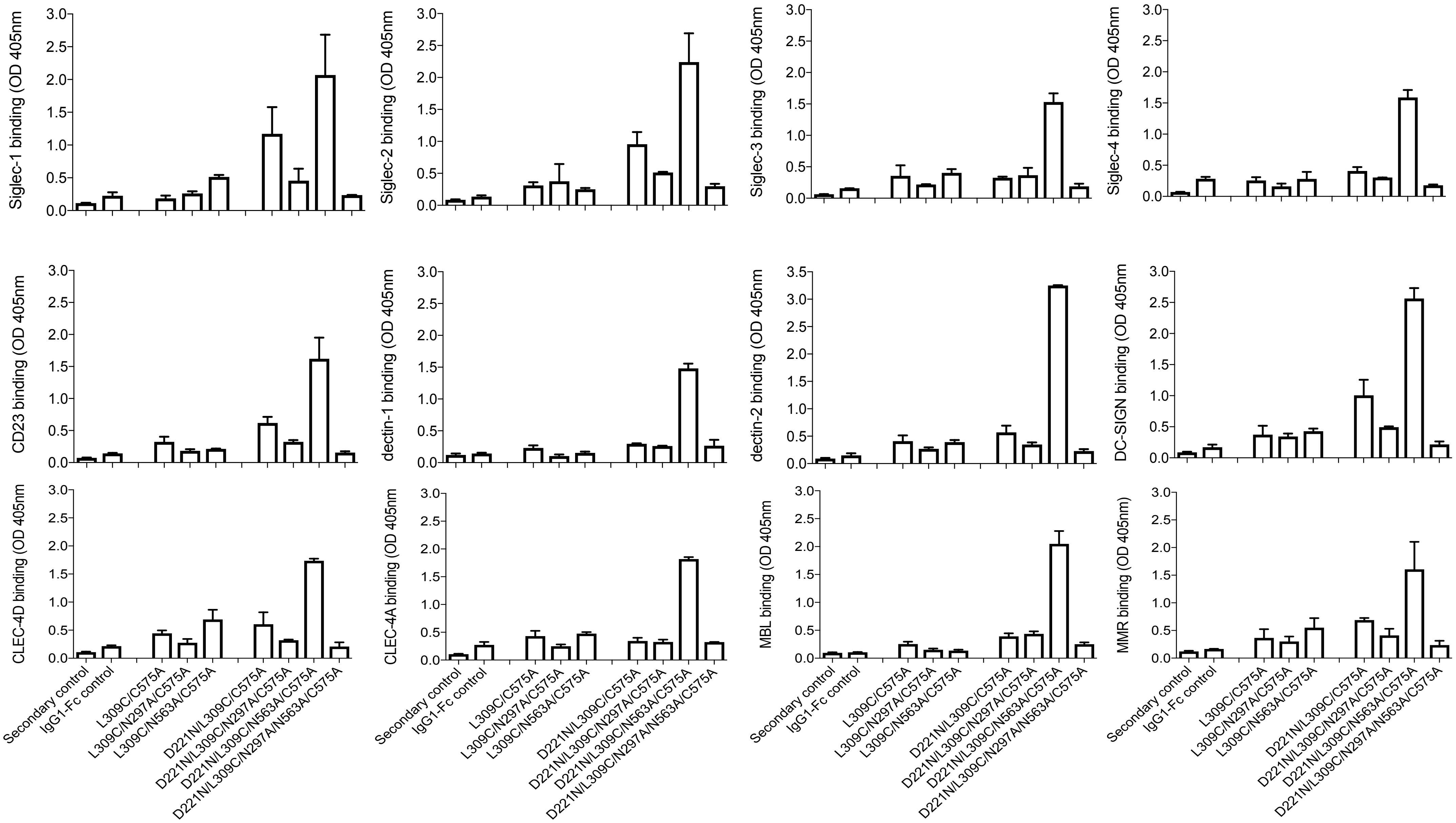

### Glycan receptor binding data for series 2 mutants

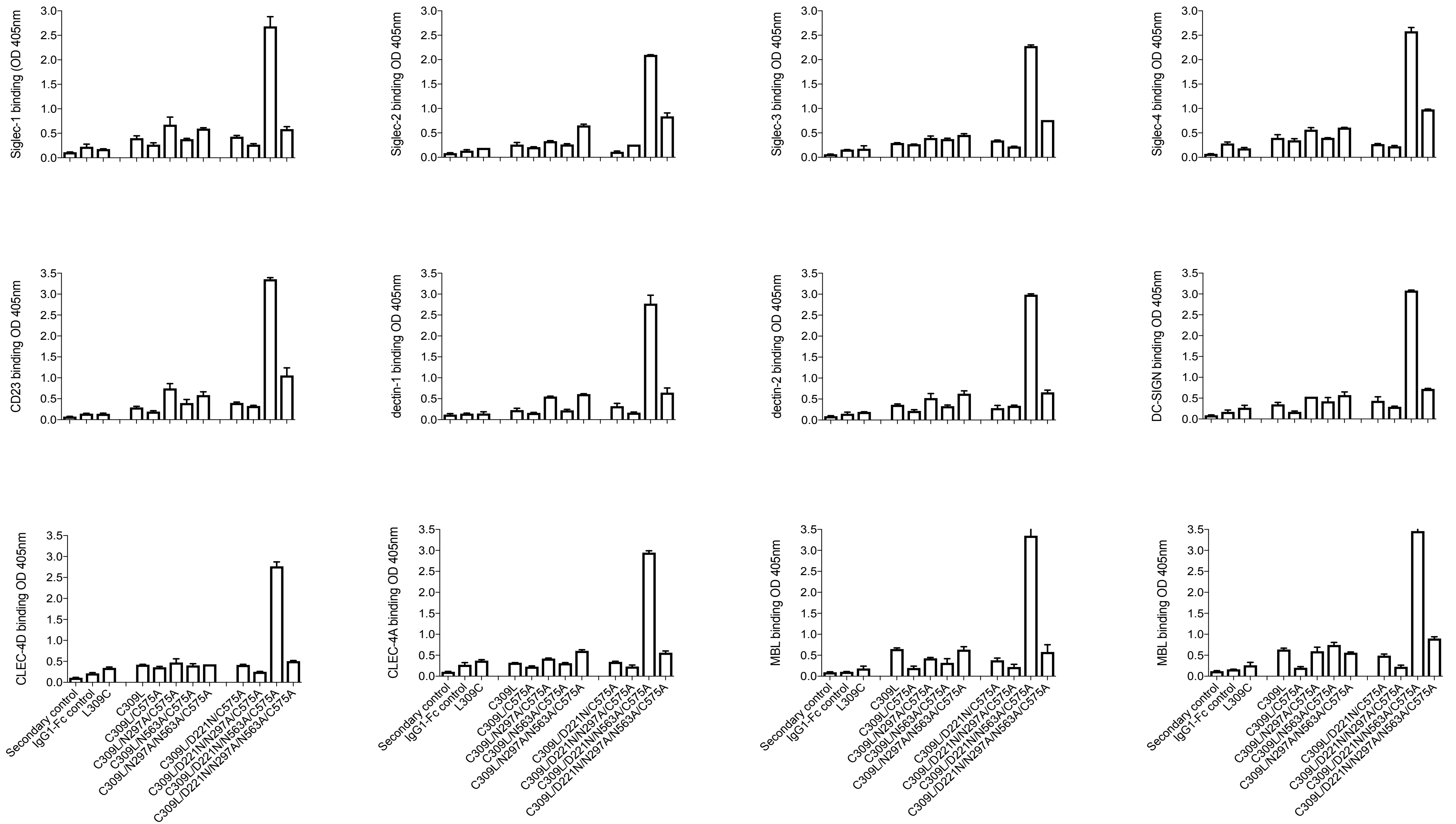

### Fcγ-receptor binding data for series 1 mutants – methods described as per manuscript

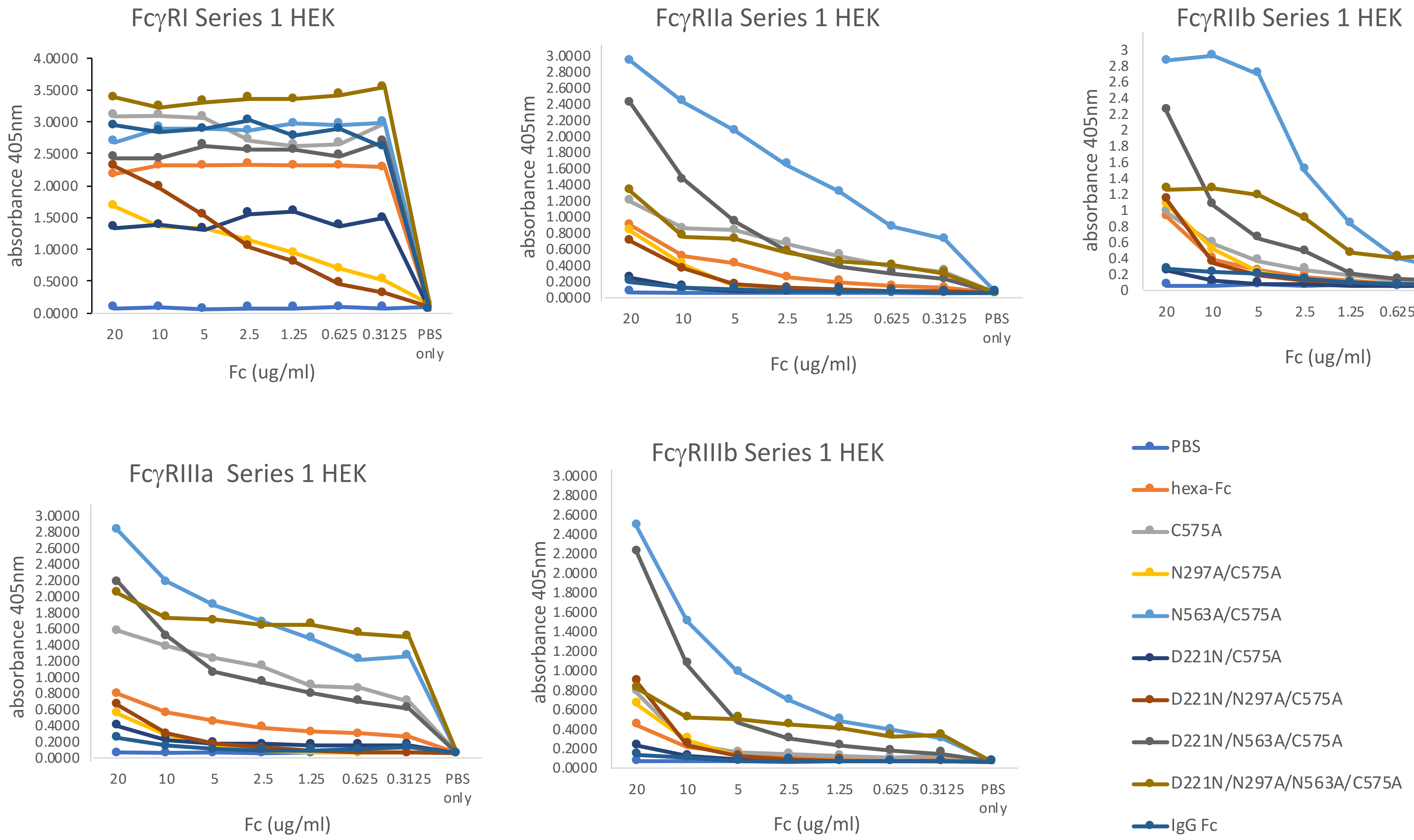

Fcγ-receptor binding data for series 2 mutants – methods described as per manuscript

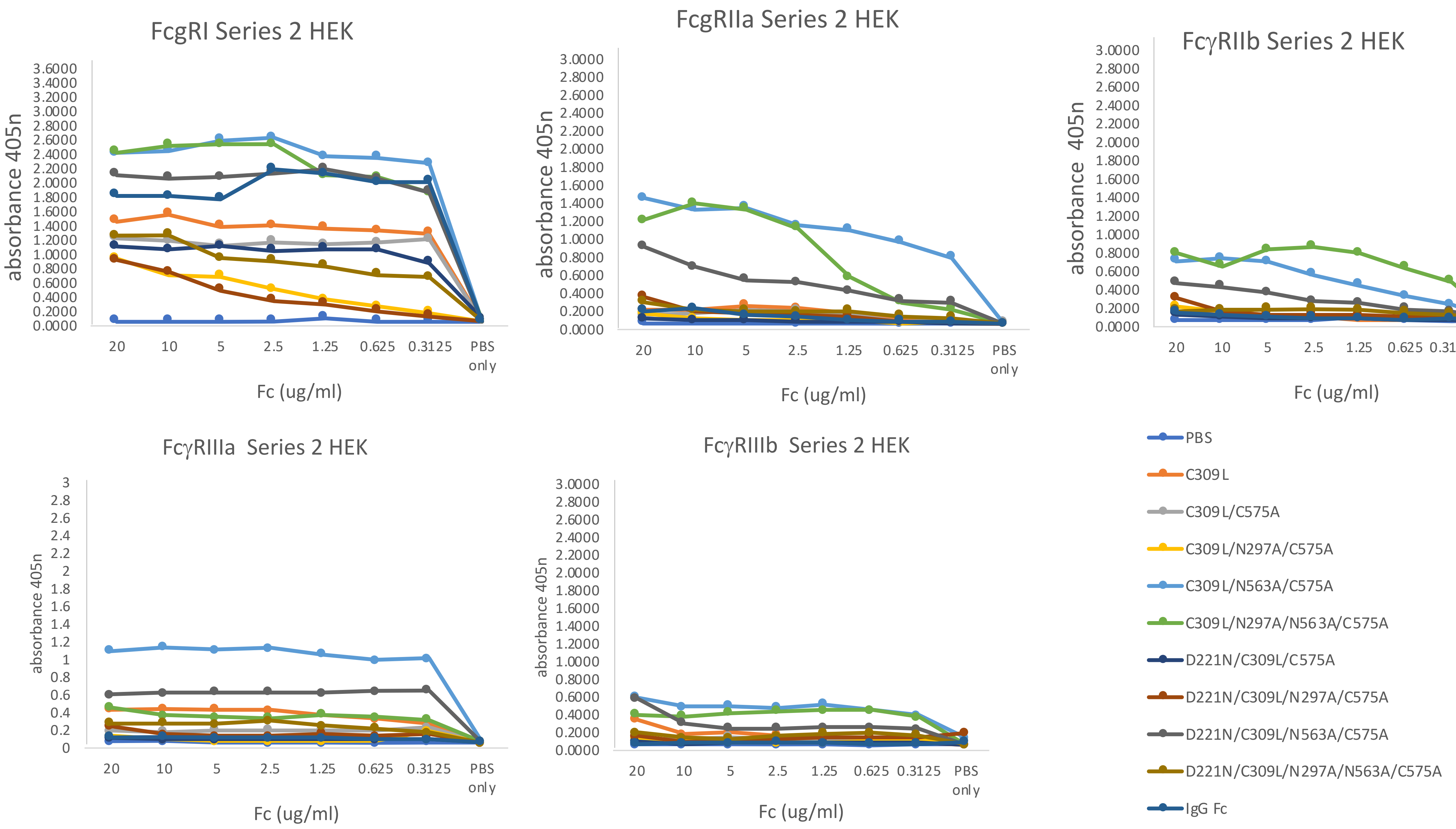

N-glycomic analysis for each mutant specified and using methods as described in manuscript

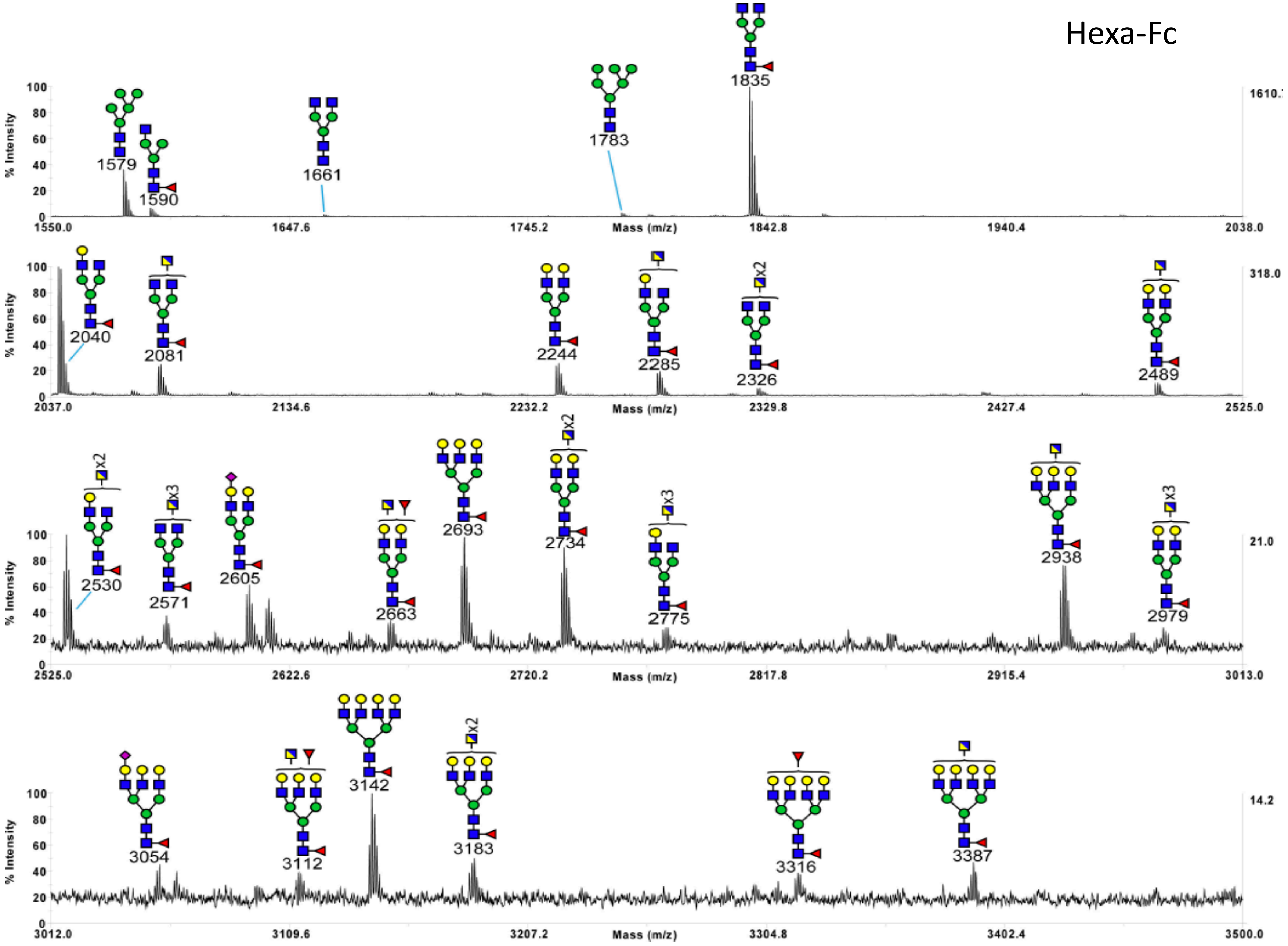

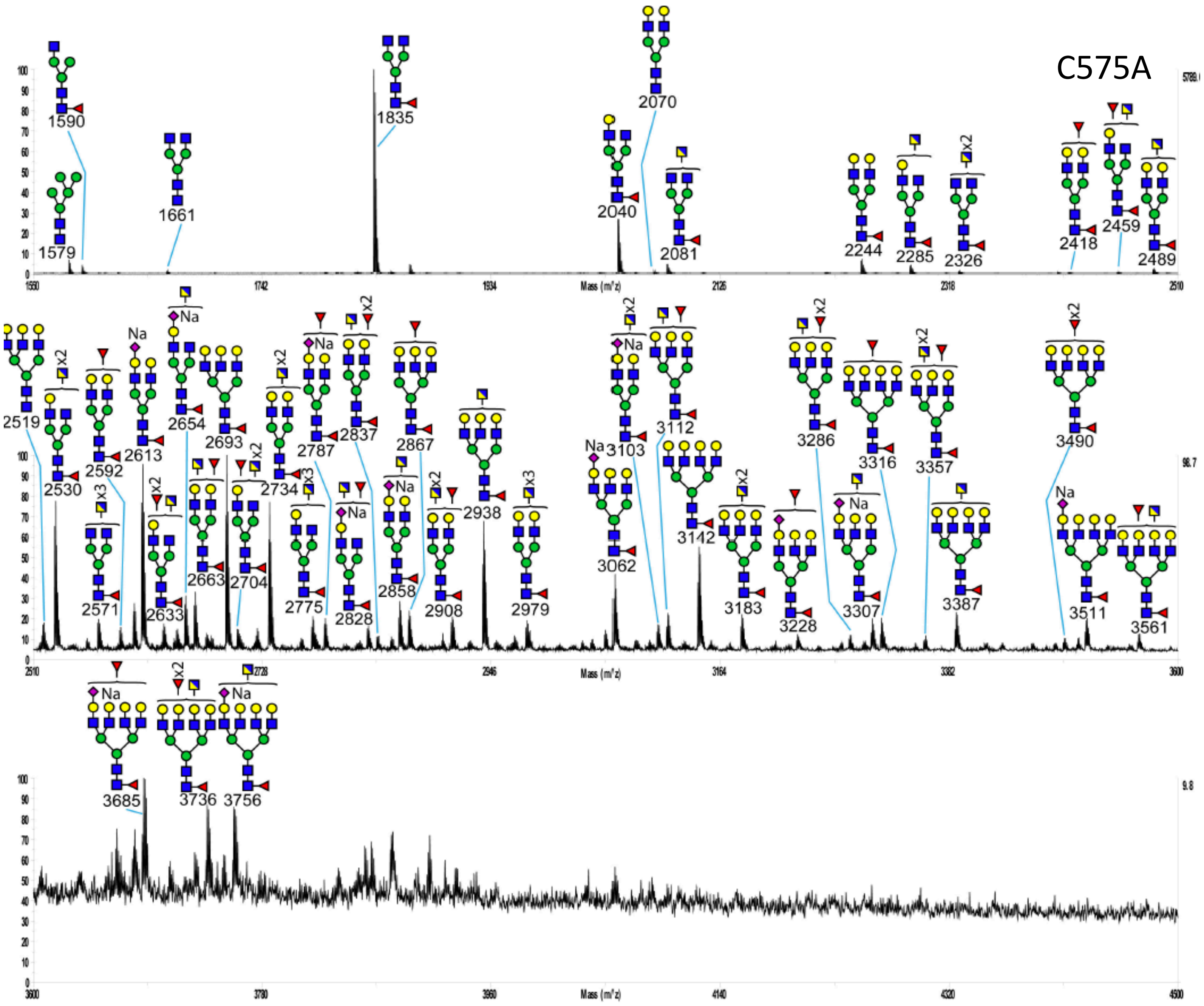

C575A
1579
1590
1661
1835
2040
2070
2081
2244
2285
2326
2418
2459
2489
Mass (m/z)
2519
2530
2571
2592
2613
2633
2654
2663
2693
2704
2734
2775
2787
2828
2837
2858
2867
2908
2938
2979
3062
3103
3112
3142
3183
3228
3286
3307
3316
3357
3387
3490
3511
3561
Mass (m/z)
3685
3736
3756
Mass (m/z)
Na
x2
x3

N297A/C575A

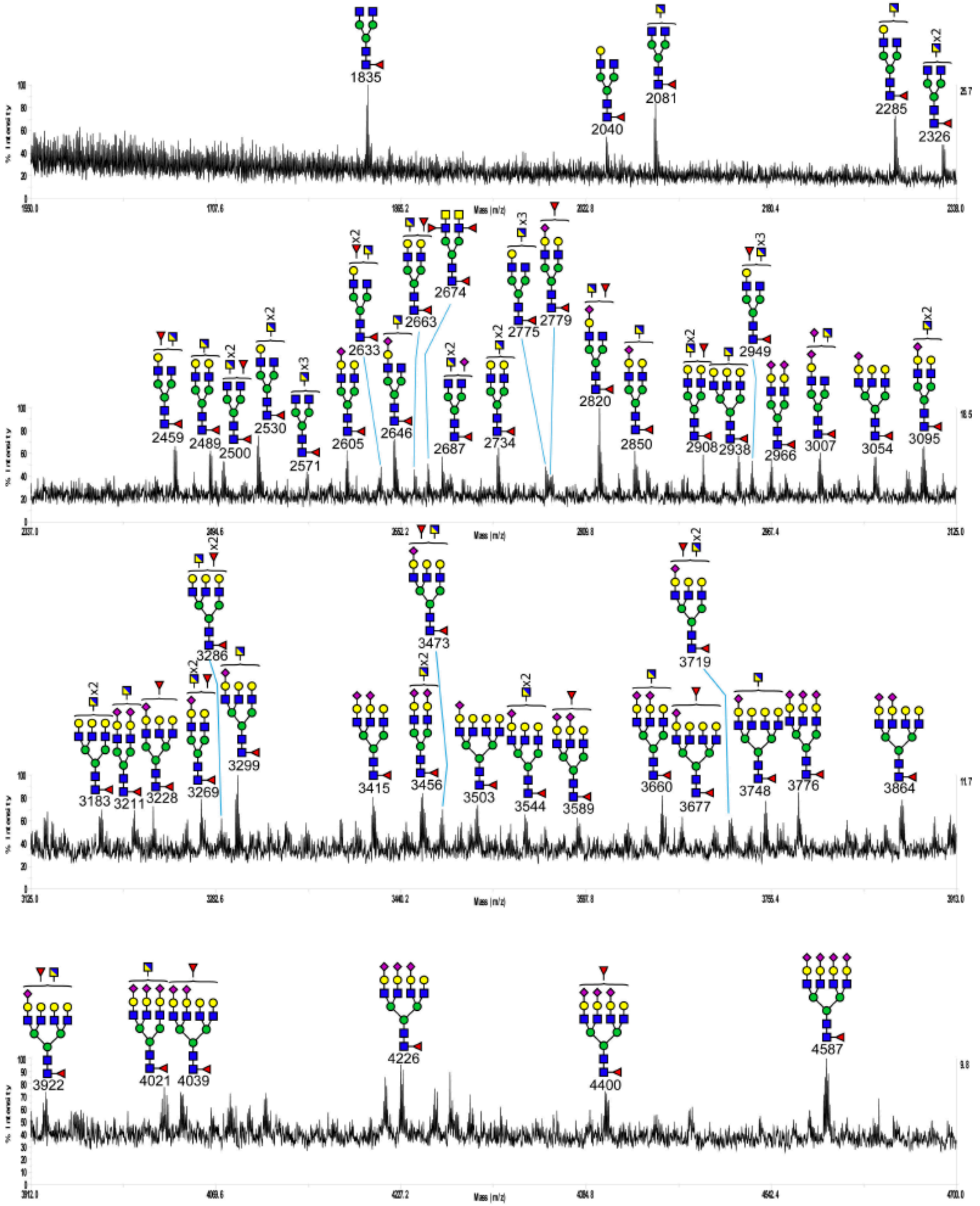
1835
2040
2081
2285
2326
2459
2489
2500
2530
2571
2605
2633
2646
2663
2674
2687
2734
2775
2779
2820
2850
2908
2938
2949
2966
3007
3054
3095
3183
3211
3228
3269
3286
3299
3415
3456
3473
3503
3544
3589
3660
3677
3719
3748
3776
3864
3922
4021
4039
4226
4400
4587
% Intensity
Mass (m/z)

N563A/C575A

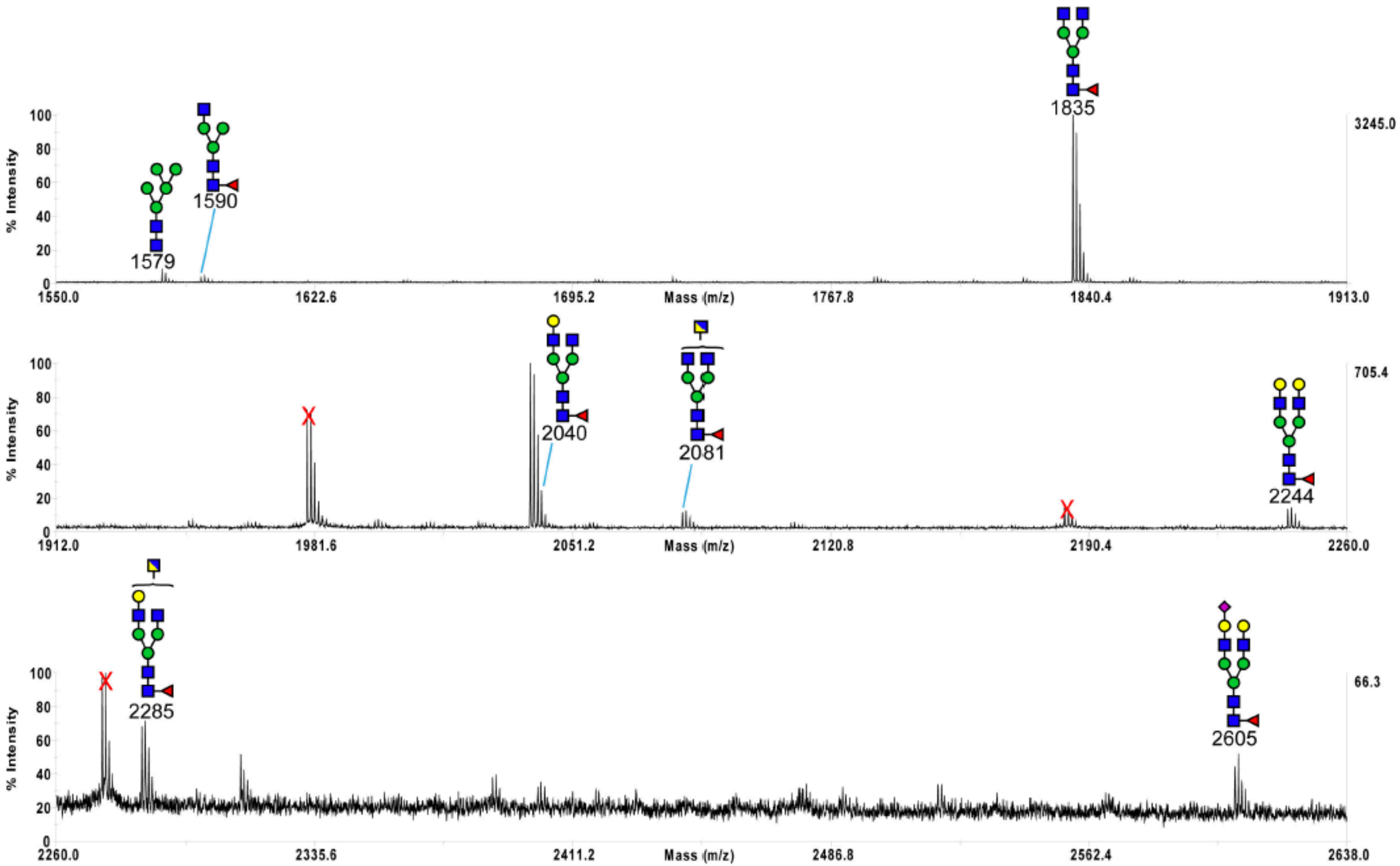

1579
1590
1835
2040
2081
2244
2285
2605
% Intensity
Mass (m/z)
3245.0
705.4
66.3
1550.0
1622.6
1695.2
1767.8
1840.4
1913.0
1912.0
1981.6
2051.2
2120.8
2190.4
2260.0
2335.6
2411.2
2486.8
2562.4
2638.0

D221N/C575A

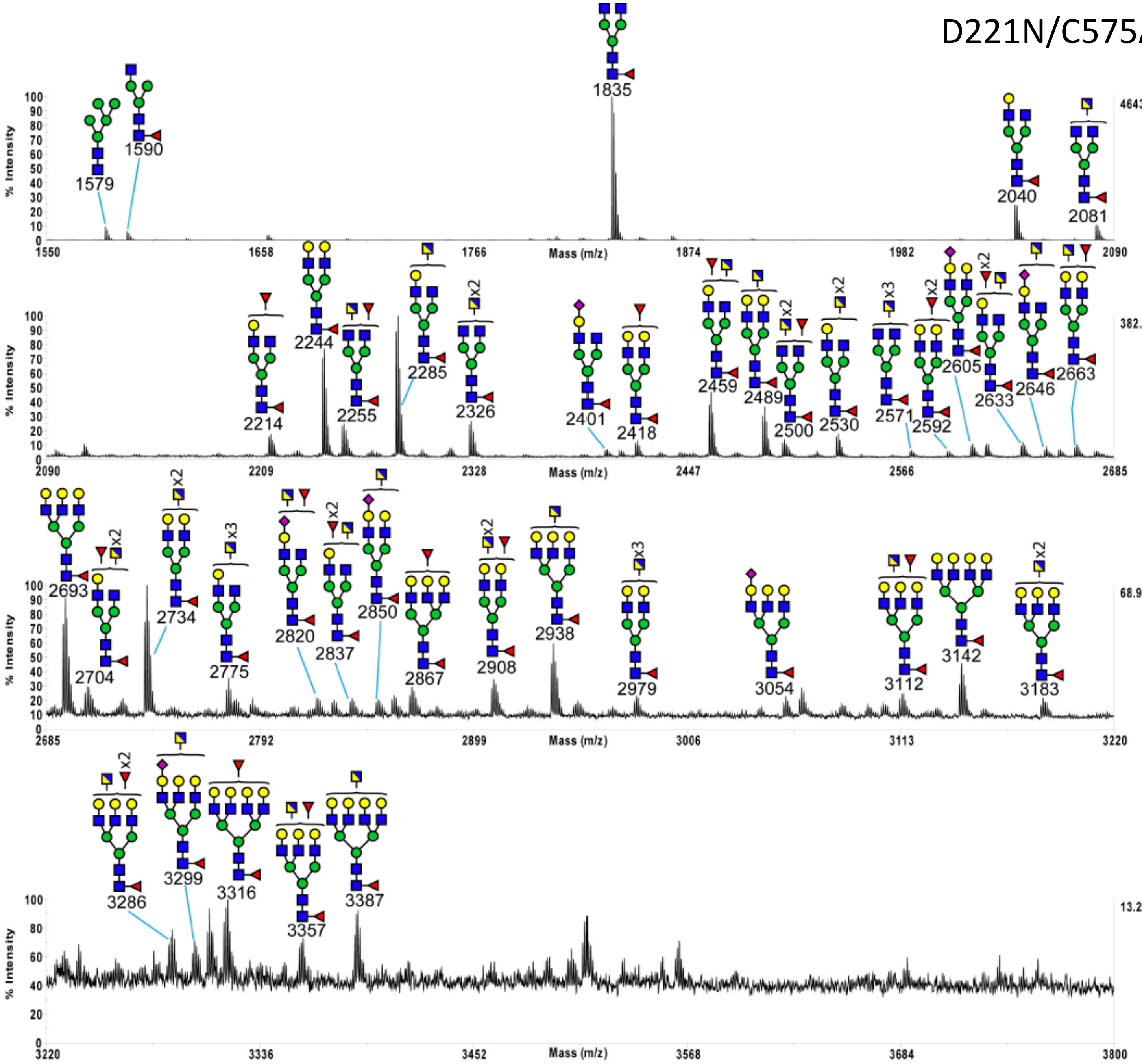

% Intensity
Mass (m/z)
4643.2
1579
1590
1835
2040
2081
1550
1658
1766
1874
1982
2090
382.3
2214
2244
2255
2285
2326
2401
2418
2459
2489
2500
2530
2571
2592
2605
2633
2646
2663
x2
x3
2209
2328
2447
2566
2685
68.9
2693
2704
2734
2775
2820
2837
2850
2867
2908
2938
2979
3054
3112
3142
3183
2792
2899
3006
3113
3220
13.2
3286
3299
3316
3357
3387
3336
3452
3568
3684
3800

D221N/N297A/C575A

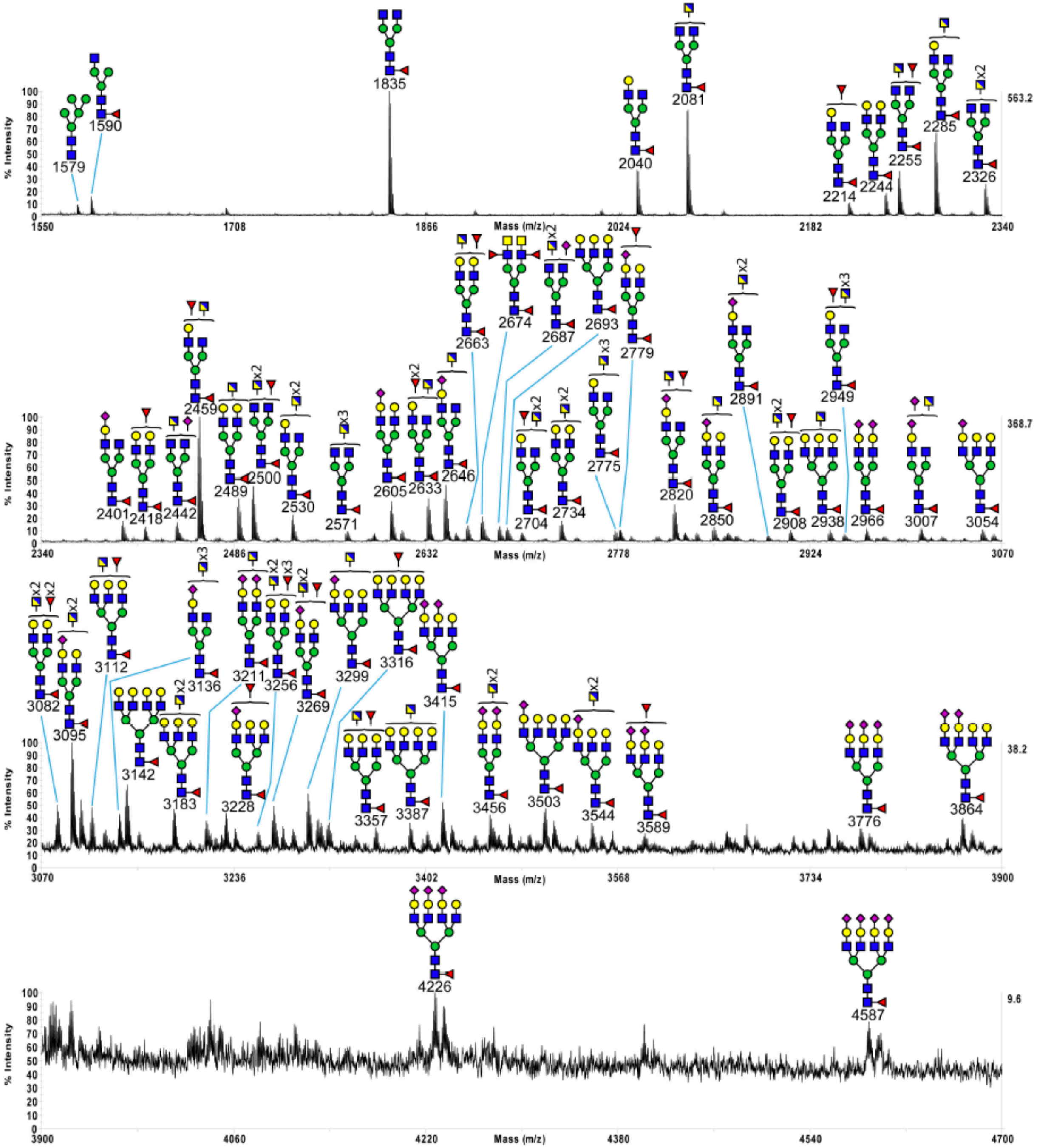

D221N/N563A/C575A

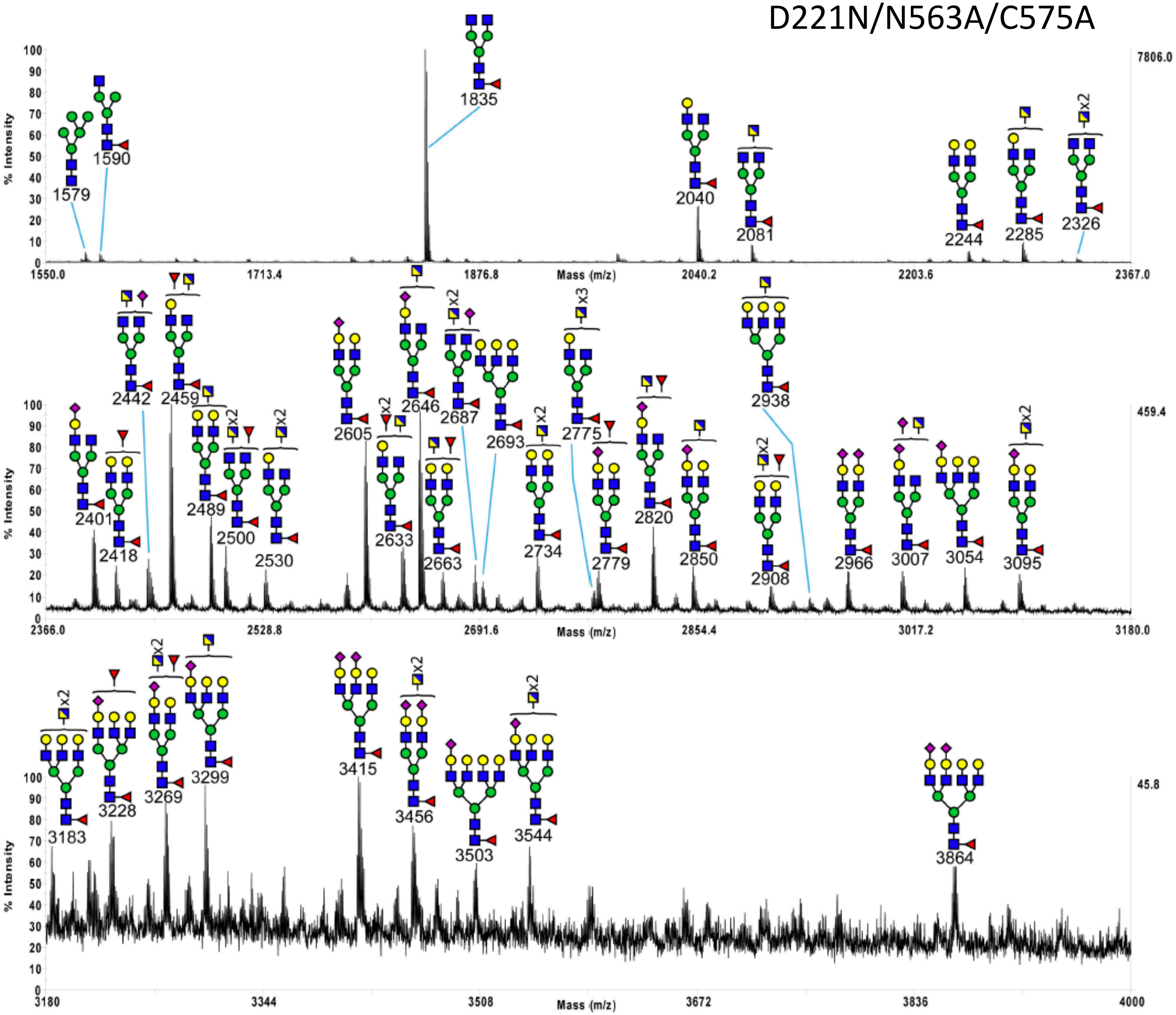

D221N/N297A/N563A/C575A

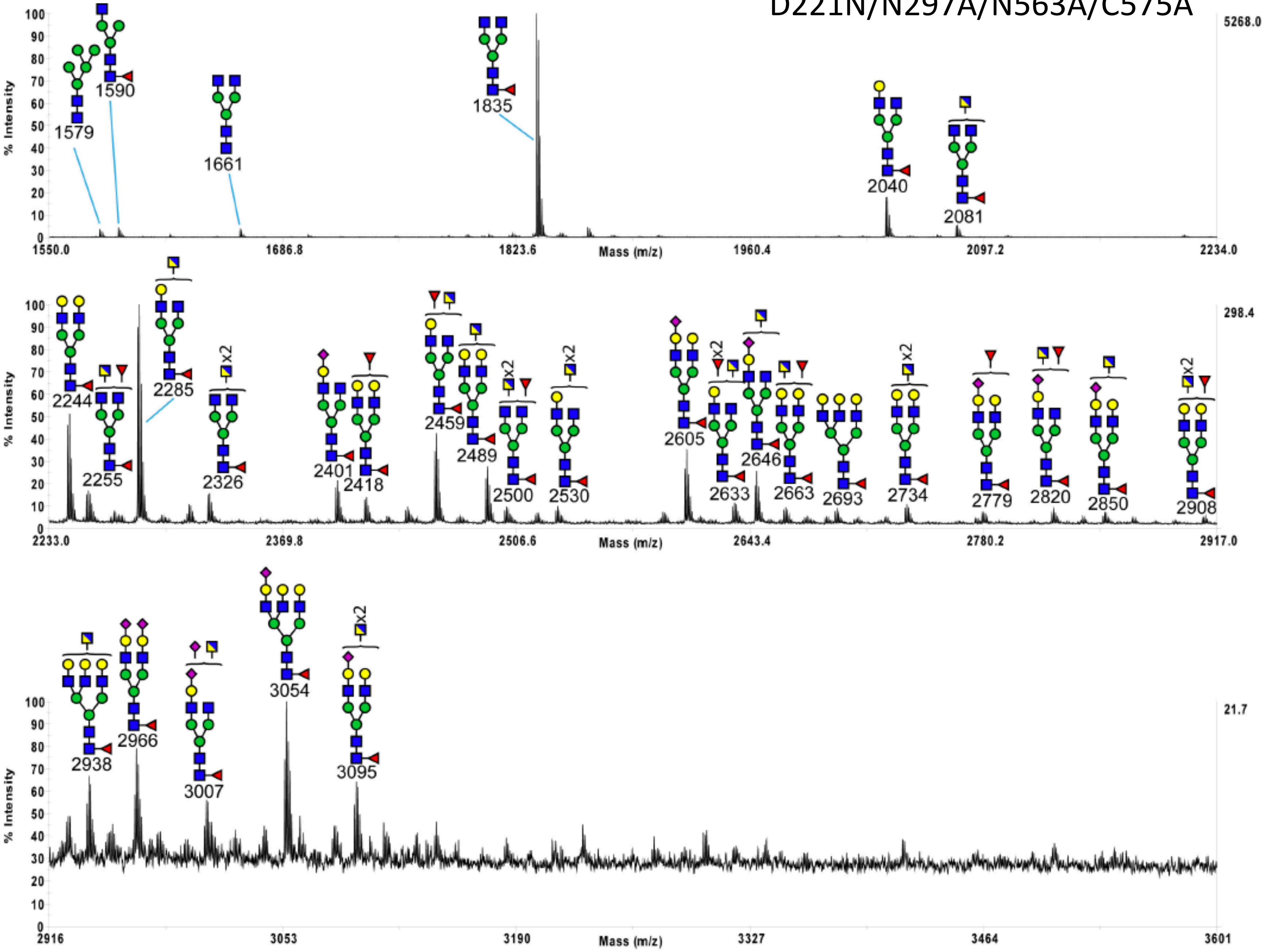

# C309L

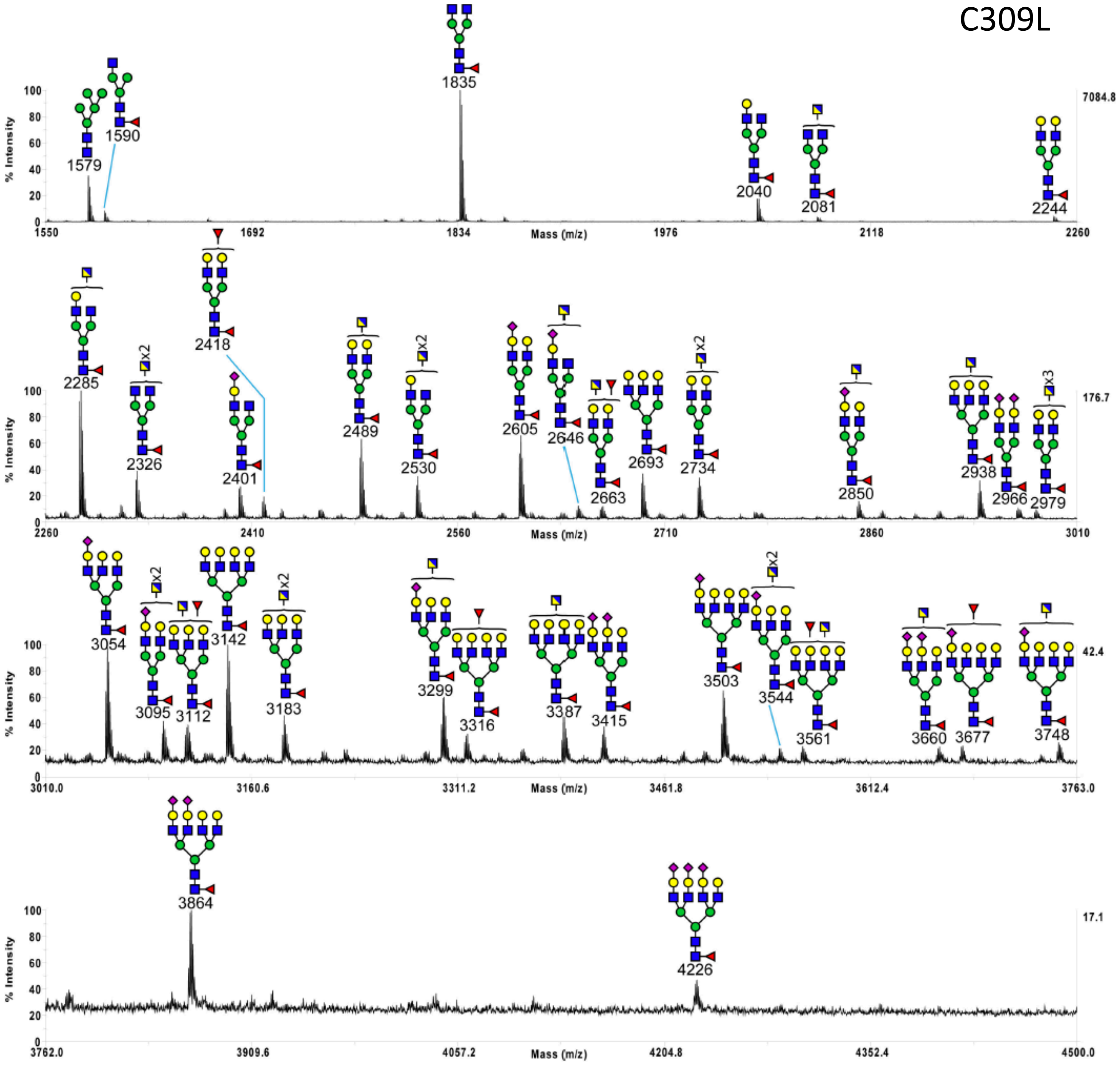
% Intensity
Mass (m/z)
7084.8
176.7
42.4
17.1
1579
1590
1835
2040
2081
2244
2285
2326
2418
2401
2489
2530
2605
2646
2663
2693
2734
2850
2938
2966
2979
3054
3095
3112
3142
3183
3299
3316
3387
3415
3503
3544
3561
3660
3677
3748
3864
4226
x2
x3
1550
1692
1834
1976
2118
2260
2410
2560
2710
2860
3010
3010.0
3160.6
3311.2
3461.8
3612.4
3763.0
3762.0
3909.6
4057.2
4204.8
4352.4
4500.0

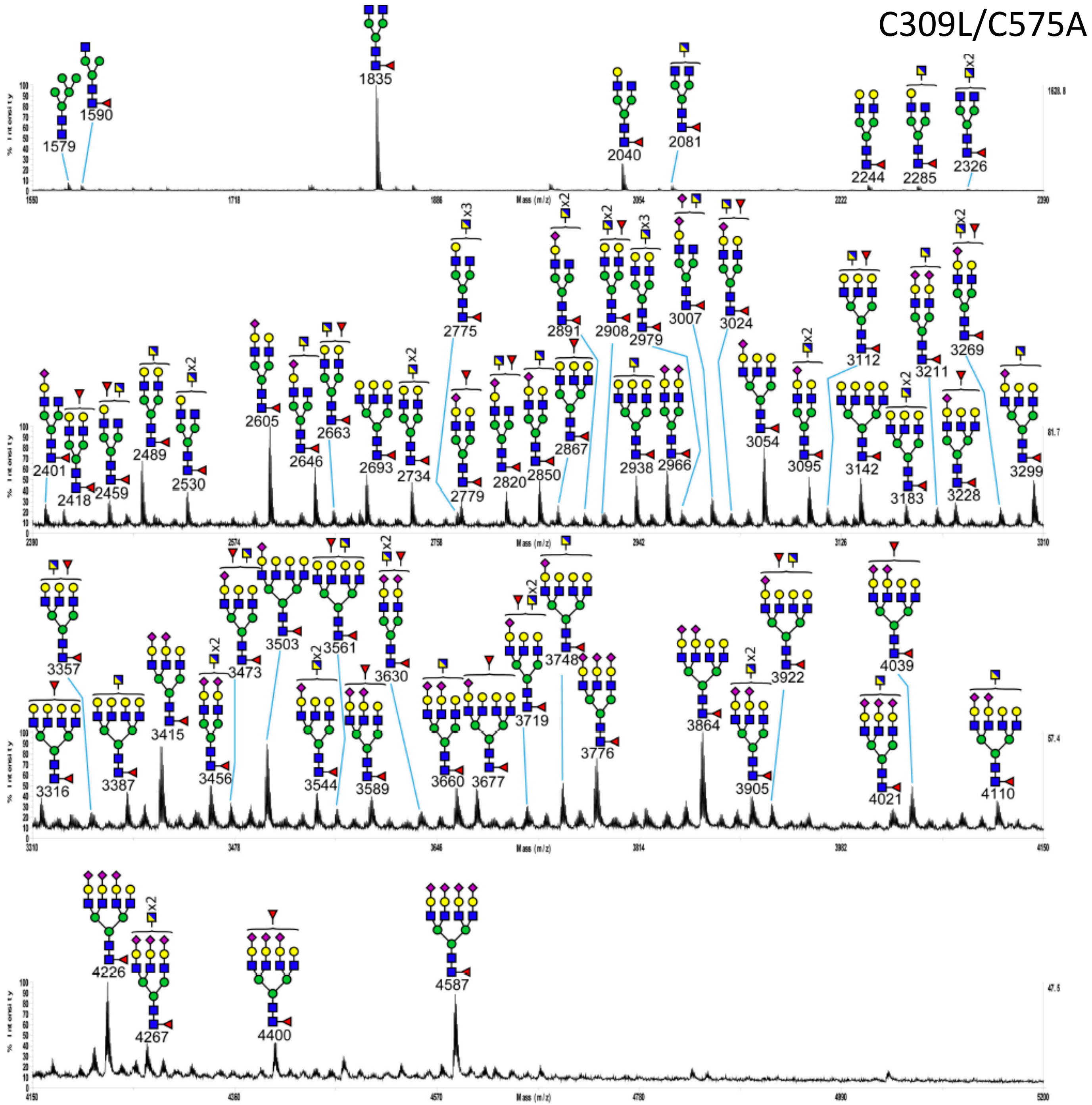
C309L/C575A
1579
1590
1835
2040
2081
2244
2285
2326
2401
2418
2459
2489
2530
2605
2646
2663
2693
2734
2775
2779
2820
2850
2867
2891
2908
2938
2966
2979
3007
3024
3054
3095
3112
3142
3183
3211
3228
3269
3299
3316
3357
3387
3415
3456
3473
3503
3544
3561
3589
3630
3660
3677
3719
3748
3776
3864
3905
3922
4021
4039
4110
4226
4267
4400
4587
% Intensity
Mass (m/z)

# C309L/N297A/C575A

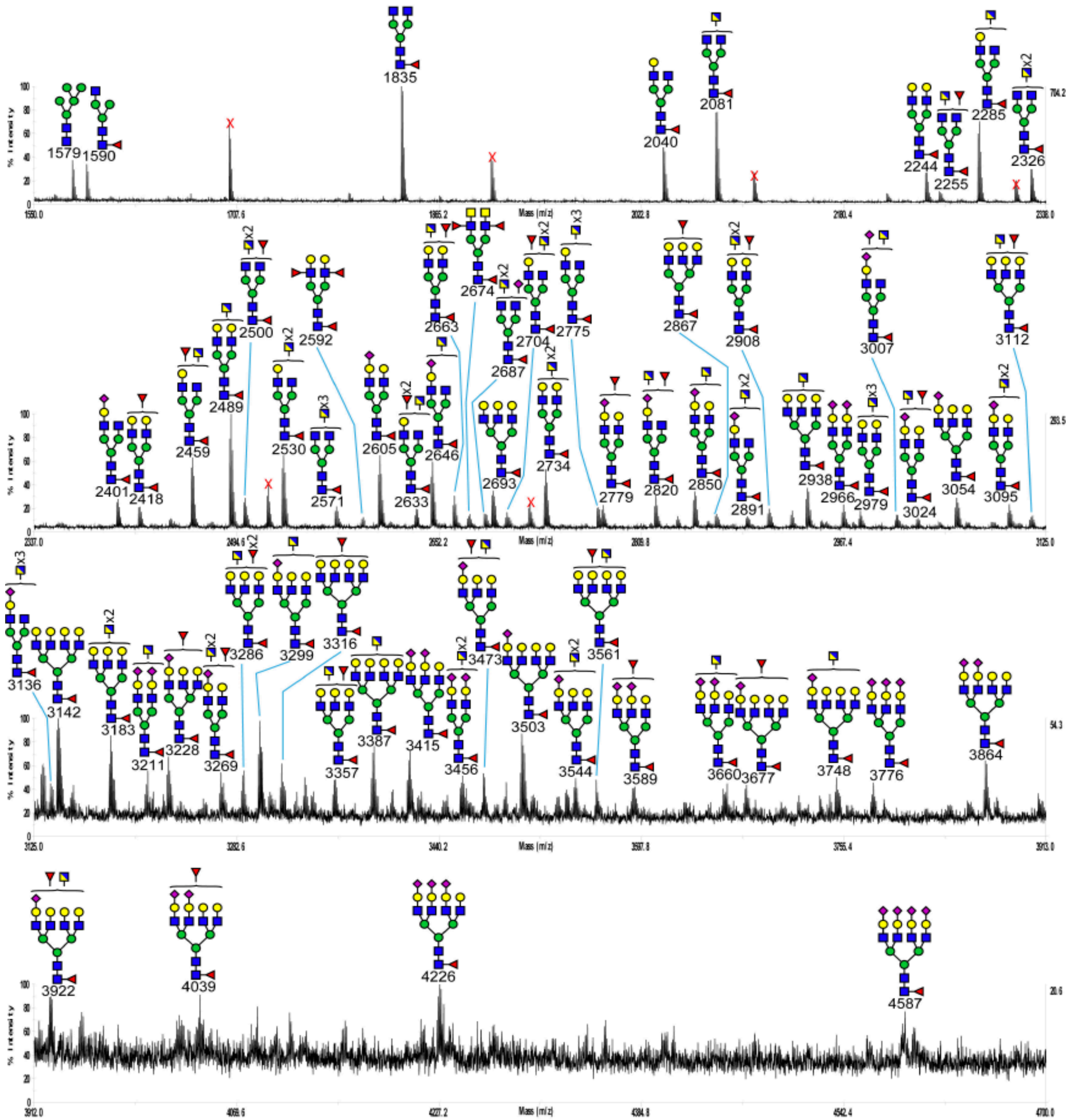

C309L/N563A/C575A

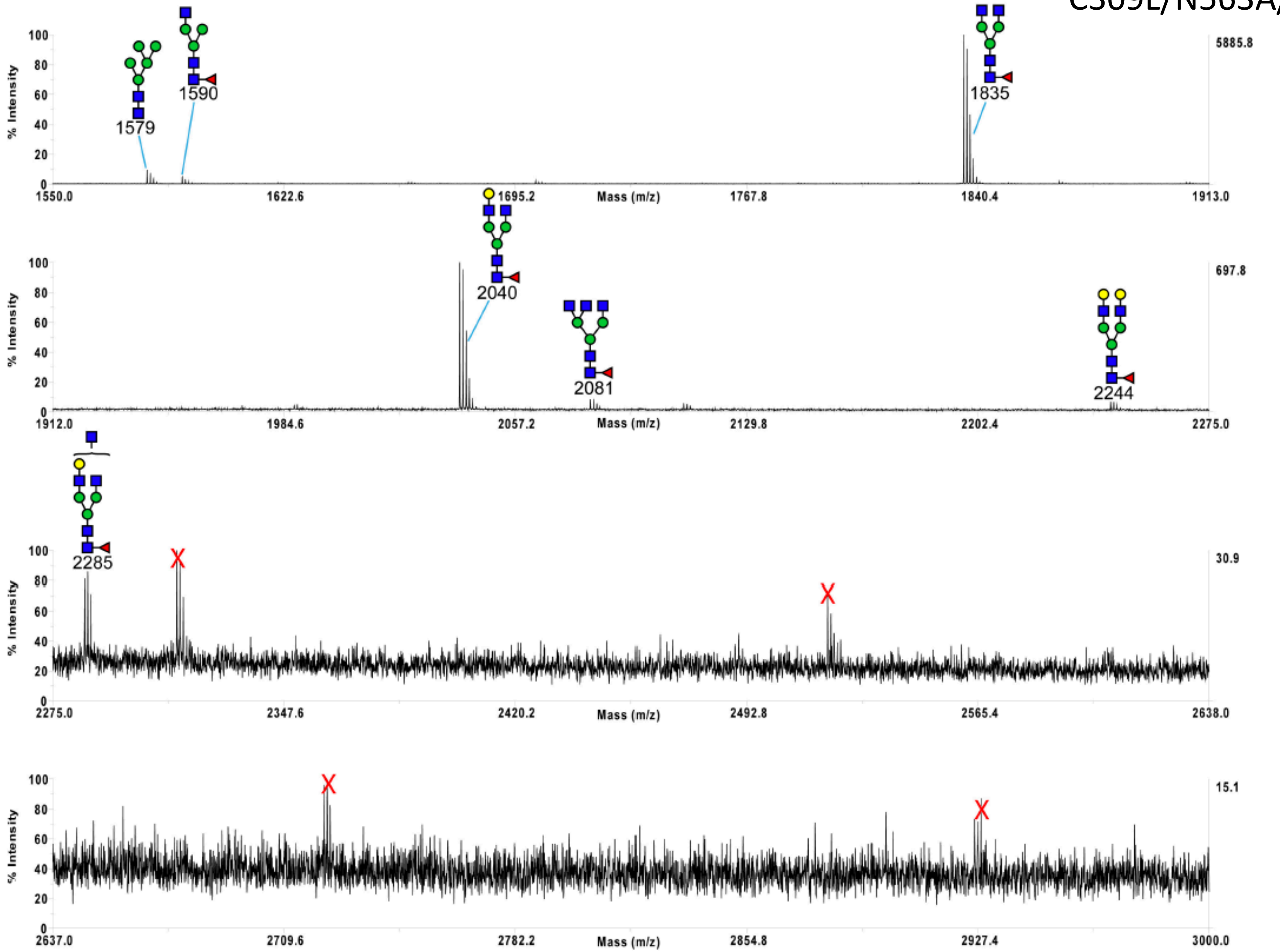

C309L/N297A/N563A/C575A

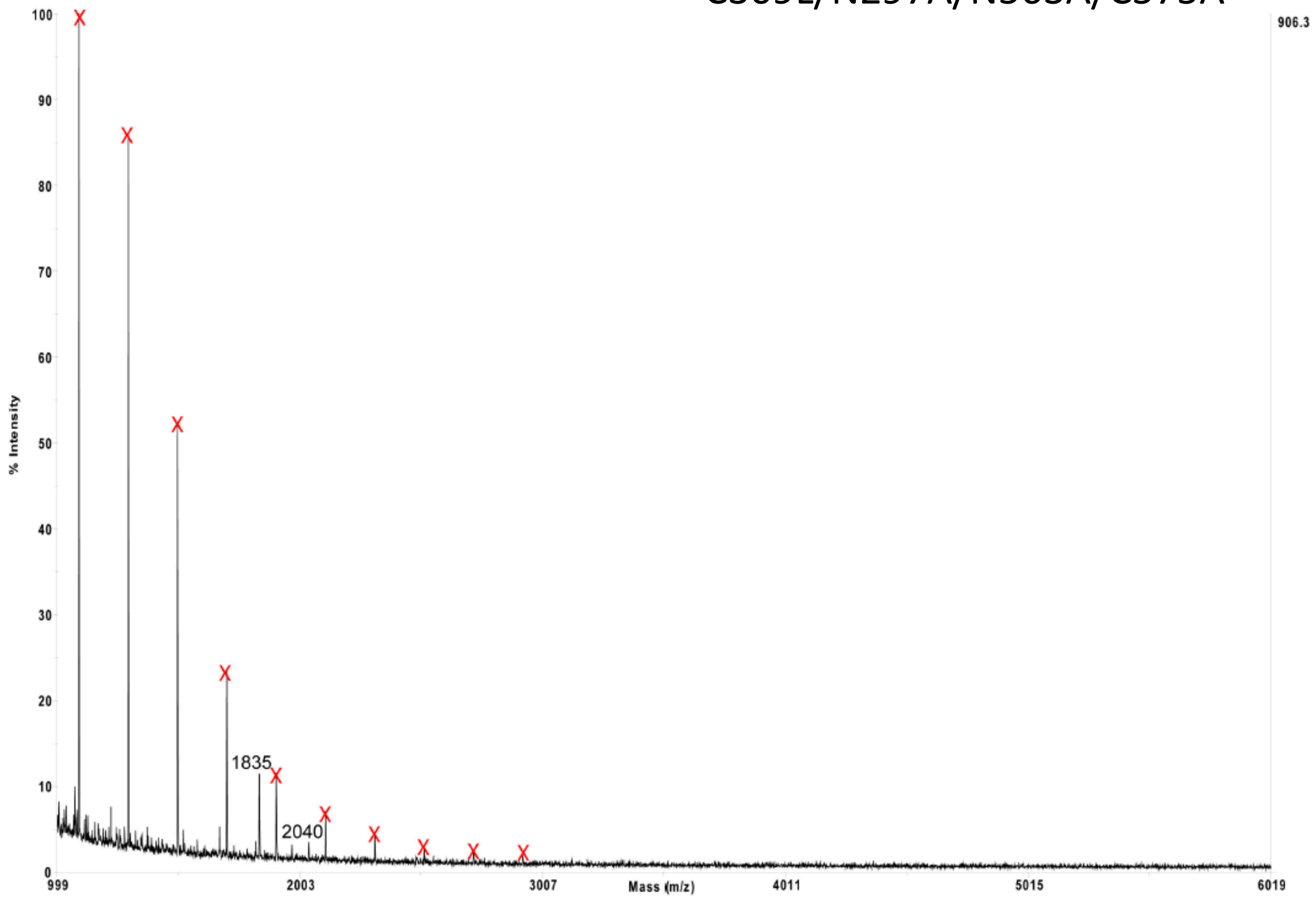

# D221N/C309L/C575A

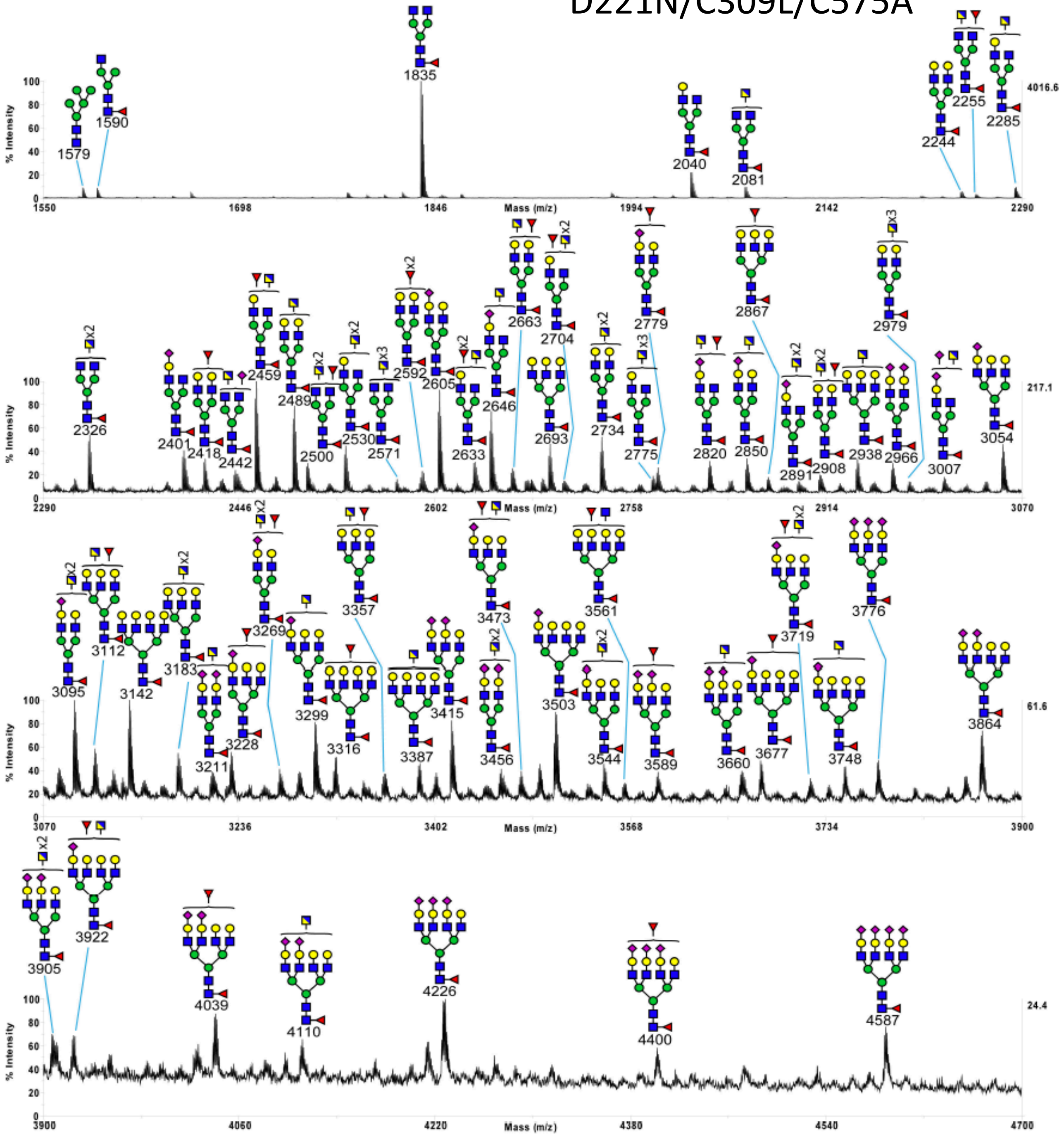

D221N/C309L/N297A/C575A

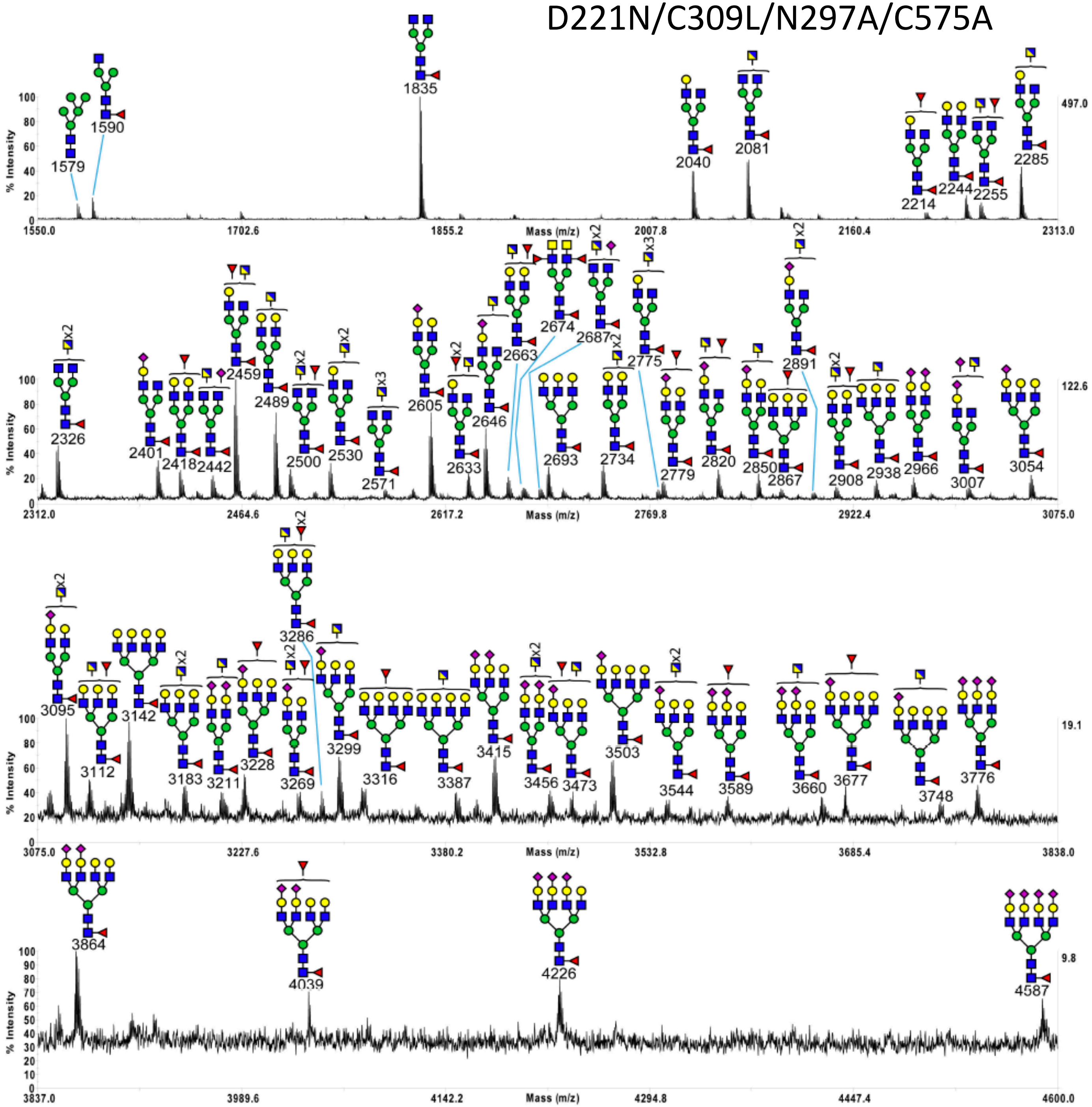

D221N/C309L/N563A/C575A

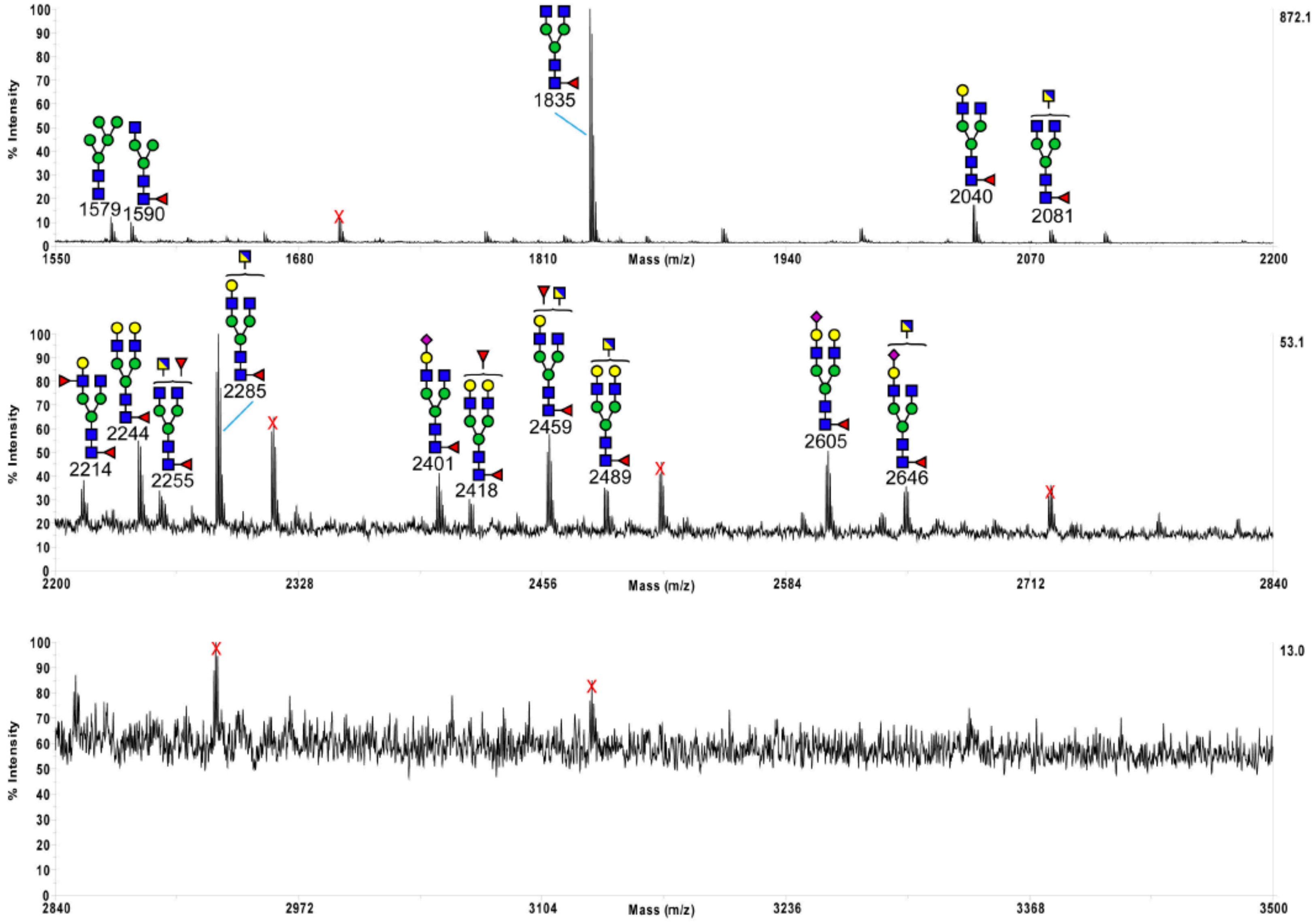

D221N/C309L/N297A/N563A
/C575A

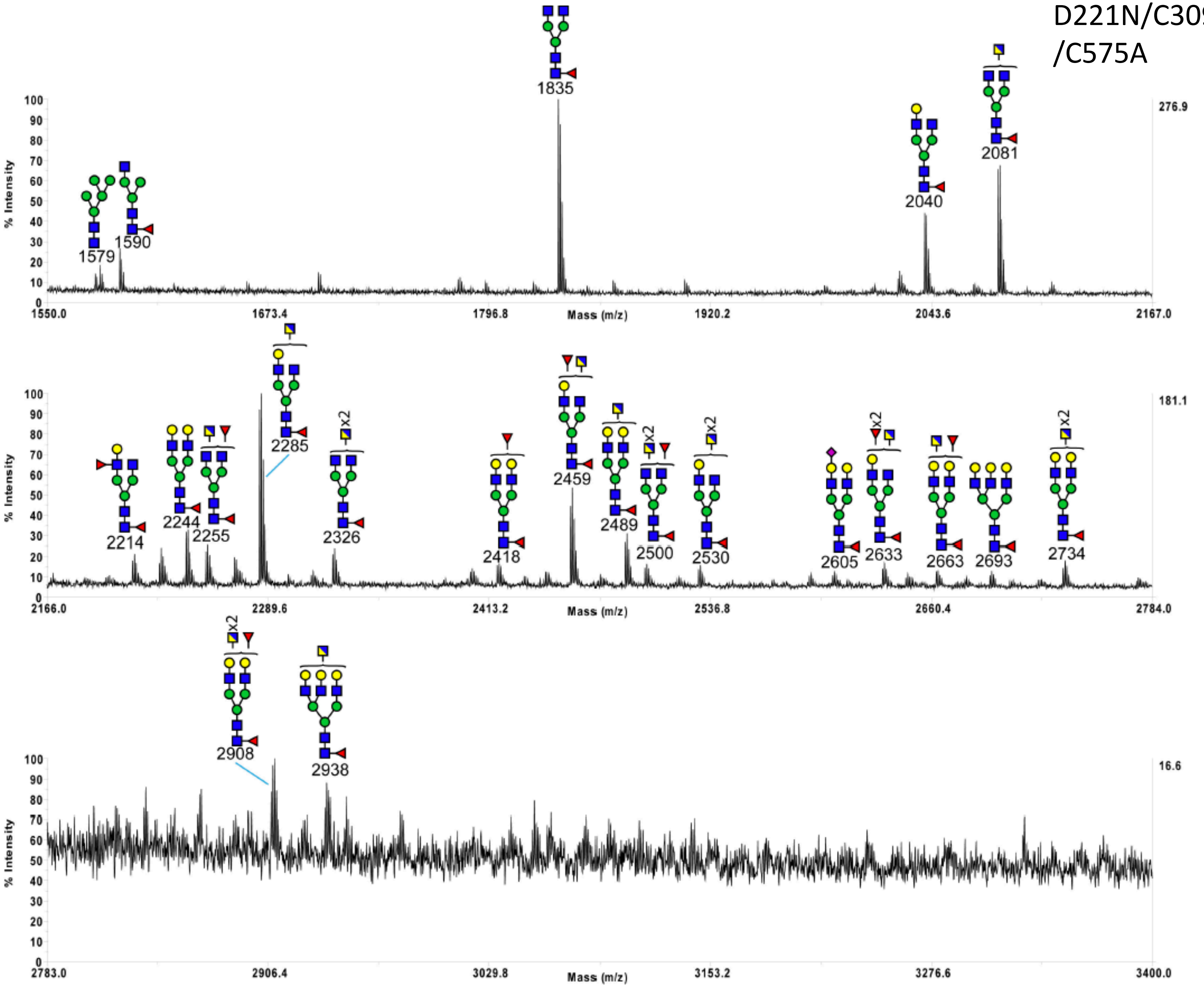
